# Eco-evolutionary dynamics of partially migratory metapopulations in spatially and seasonally varying environments

**DOI:** 10.1101/2023.11.28.568986

**Authors:** Thomas R. Haaland, Ana Payo-Payo, Paul Acker, Rita Fortuna, Sarah J. Burthe, Irja I. Ratikainen, Francis Daunt, Jane M. Reid

**Affiliations:** Department of Biology, Norwegian University of Science and Technology, Trondheim, Norway; Departamento de Biodiversidad, Ecología y Evolución, Universidad Complutense de Madrid, Madrid, Spain; UK Centre for Ecology & Hydrology, Bush Estate, Penicuik, United Kingdom; School of Biological Sciences, University of Aberdeen, United Kingdom

**Keywords:** spatial ecology, seasonal migration, threshold trait, phenotypic polymorphism, environmental stochasticity, liability, alternative tactics, extreme climatic events, life-history trait, partial migration, meta-population, quantitative trait, evolutionary rescue

## Abstract

Predicting population responses to environmental changes requires understanding interactions among environmentally induced phenotypic variation, selection, demography and genetic variation, and thereby predicting eco-evolutionary dynamics emerging across diverse temporal and spatial scales. Partially migratory metapopulations (PMMPs), featuring seasonal coexistence of resident and migrant individuals across multiple spatially distinct subpopulations, have clear potential for complex spatio-seasonal eco-evolutionary dynamics through impacts of selection on migration on spatial population dynamics, and feedbacks resulting from ongoing micro-evolution. However, the key genetic and environmental conditions that maintain migratory polymorphisms, and eco-evolutionary dynamics of PMMPs under stochastic environmental variation and strong seasonal perturbations, have not yet been resolved. Accordingly, we present a general individual-based model that tracks eco-evolutionary dynamics in PMMPs inhabiting spatially structured, seasonally varying landscapes, with migration formulated as a quantitative genetic threshold trait. Our simulations show that such genetic and landscape structures, which commonly occur in nature, can readily produce a variety of stable partially migratory systems given diverse regimes of spatio-seasonal environmental variation. Typically, partial migration is maintained whenever sites differ in non-breeding season suitability resulting from variation in density-dependence, causing ‘ideal free’ non-breeding distributions where residents and migrants occur with frequencies that generate similar survival probabilities. Yet, stable partial migration can also arise without any fixed differences in non-breeding season density-dependence among sites, and even without density-dependence at all, through risk-spreading given sufficiently large stochastic environmental fluctuations among sites and years. Finally, we show that local non-breeding season mortality events, as could result from extreme climatic events, can generate eco-evolutionary dynamics that ripple out to affect breeding and non-breeding season space use of subpopulations throughout the PMMP, on both short and longer timeframes. Such effects result from spatially divergent selection on both the occurrence and destinations of migration. Our model thus shows how facultative seasonal migration can act as a key mediator of eco-evolutionary dynamics in spatially and seasonally structured environments, providing key steps towards predicting responses of natural partially migratory populations to ongoing changes in spatio-seasonal patterns of environmental variation.

## Introduction

It is increasingly clear that natural populations experience ecological and evolutionary changes on comparable timeframes, generating complex eco-evolutionary outcomes (Pelletier et al. 2007, Ellner et al. 2011, Lion 2018, Govaert et al. 2019). Further, numerous populations are experiencing rapid human-induced environmental changes, including increasing frequencies and severities of extreme climatic events (ECEs) such as droughts, heatwaves, floods and storms (Hirabayashi et al. 2013, Trenberth et al. 2015, Stott 2016). Such disturbances can impact vital rates, causing immediate population declines coupled with episodes of strong natural selection (Van de Pol et al. 2017). Meanwhile, theoretical and empirical studies indicate that the potential for eco-evolutionary feedbacks is greatest when selection is strong (Reznick and Ghalambor 2001, Lavergne et al. 2010, terHorst and Zee 2016). Integrated understanding of forms and mechanisms of eco-evolutionary dynamics induced by diverse regimes of environmental variation, including ECEs, is therefore required to predict and manage population outcomes (Kinnison and Hairston 2007, Hanski 2012, Ferriere and Legendre 2013).

Rapid eco-evolutionary dynamics fundamentally require the presence of phenotypic and genetic variation in fitness-related traits. Systems that exhibit life-history polymorphisms are therefore particularly pertinent for studying how novel environmental variation can generate eco-evolutionary responses occurring across diverse spatial and temporal scales (Bell and Collins 2008, Lawson et al. 2015). Such work could also address the foundational questions of how phenotypic polymorphisms, and underlying genetic variation, are maintained.

One striking life-history polymorphism concerns partial migration, for example where non-migratory individuals that remain resident in their breeding area all year seasonally coexist with migrants that leave during non-breeding seasons then return to breed (Chapman et al. 2011). Such partial migration is taxonomically and geographically widespread, occurring in numerous birds, reptiles, fish, amphibians and mammals, and involving short- or long-distance seasonal movements (Grayson et al. 2011, Dodson et al. 2013, Berg et al. 2019, Chambon et al. 2019). Spatial variation in seasonal environmental conditions can then induce strong selection on residence versus migration, through heritable components of migratory strategy and/or carry-over effects on survival and/or subsequent reproduction (Reid et al. 2018, Ohms et al. 2019, Acker et al. 2021a). Further, the degree and form of migration directly determines seasonal distributions of individuals, shaping spatio-seasonal population dynamics (Kasai et al. 2018; Martin et al. 2022). Partially migratory systems consequently have substantial intrinsic potential for complex spatio-seasonal eco-evolutionary dynamics. For example, a local ECE could cause strong selection for or against migration, with resulting micro-evolution altering subsequent frequencies of migrants, and hence spatio-seasonal population dynamics. Density-dependence in vital rates can then cause frequency-dependence in the fitness consequences of residence versus migration, further affecting micro-evolutionary outcomes (Kaitala et al. 1993, Taylor and Norris 2007). Overall population and evolutionary dynamics could consequently play out over spatial and temporal scales far exceeding the original ECE (Reid et al. 2018). Yet, such spatio-seasonal eco-evolutionary dynamics, and their implications for the maintenance of migratory polymorphisms and overall (meta)population viability, remain largely unexplored.

Existing partial migration theory primarily envisages simple two-site systems where one site can be occupied all year and the other site is suitable in only one season, and simple genetic architectures such as a single locus with competing ‘resident’ and ‘migrant’ alleles (Taylor and Norris 2007, Griswold et al. 2010, Kokko 2011), although both spatial and genetic model variations exist (Taylor and Norris 2010, Reid et al. 2018, Ohms et al. 2019). Overall, such models show that stable polymorphisms can be maintained given balanced (i.e. equal) fitness of residents and migrants, and/or negative density-dependence at one or both sites generating negative frequency-dependent selection (Lundberg 1987; Taylor and Norris 2007; Shaw and Levin 2011; Ohms et al. 2019; Kokko and Lundberg 2001; Kaitala et al. 1993; Griswold et al. 2011). Thus, partial migration is predicted to be maintained only under rather narrow conditions, defined by strengths and forms of density-dependence, and balanced fitness effects. However, such predictions are not congruent with observations that partial migration is prevalent across highly diverse conditions in nature (Chapman et al. 2011). Although data are still scarce, a meta-analysis comparing fitness components of alternative migratory tactics in 18 animal species found that effects were approximately balanced only in 5% of comparisons (Buchan et al. 2020). New models of partial migration that relax current restrictive assumptions concerning spatial structures and genetic variation could provide new insights, by revealing wider conditions that foster polymorphisms, and revealing the resulting potential for spatio-seasonal eco-evolutionary dynamics.

First, we should relax the highly restrictive assumptions that partial migration only involves two sites, and that migrants move into empty space. In nature, migrants from one population can arrive at sites holding year-round residents and/or migrants from other populations (Fig. 1a). Such ‘partially migratory metapopulations’ (PMMPs, Reid et al. 2018) are likely commonplace and can encompass both ‘shared breeding’ and ‘shared non-breeding’ partial migration (Griswold et al. 2010) among seasonally interconnected subpopulations (e.g. whales, Geijer et al. 2016; ungulates, Berg et al. 2019; anadromous fish, Austin et al. 2019; flamingos, Sanz-Aguilar et al. 2012; newts, Grayson et al. 2011; songbirds, Zúñiga et al. 2017). These PMMP networks, coupled with spatio-seasonal variation in environmental conditions, may render stable partial migration typical rather than exceptional (Reid et al. 2018). For example, if a site hosts both local residents and incoming migrants from other sites during the non-breeding season, a major non-breeding season mortality event (e.g. due to extreme weather, disease, or other stressors) would cause selective removal of residents in the local subpopulation (i.e. individuals that breed in this site), resulting in strong selection for migration. Meanwhile, subpopulations from which any incoming migrants originated will simultaneously experience selection against migration. A single local perturbation thereby induces strong spatially-divergent selection. Such stochastic spatio-seasonal environmental conditions could also favour partial migration through genotype-level bet-hedging. Here, if parents can spread risk by producing offspring that use diverse non-breeding season locations (some residents; some migrants to different destinations), variance in fitness is reduced, facilitating lineage survival (Cohen 1967, but see Lundberg 1987). PMMP structures will also alter key relationships between migration frequency and seasonal population densities, thereby altering forms and magnitudes of frequency-dependent selection on migration. Yet, while previous theory highlighted how metapopulation dynamics are complicated by the existence of migratory networks (Taylor and Norris 2010, Taylor and Hall 2012, Payo-Payo et al. 2022), no work has examined spatio-seasonal eco-evolutionary dynamics in stochastic seasonal environments, including responses to ECEs to which migratory populations may be particularly vulnerable (Runge et al. 2015, Shaw 2016, Kubelka et al. 2022).

**Figure 1:**
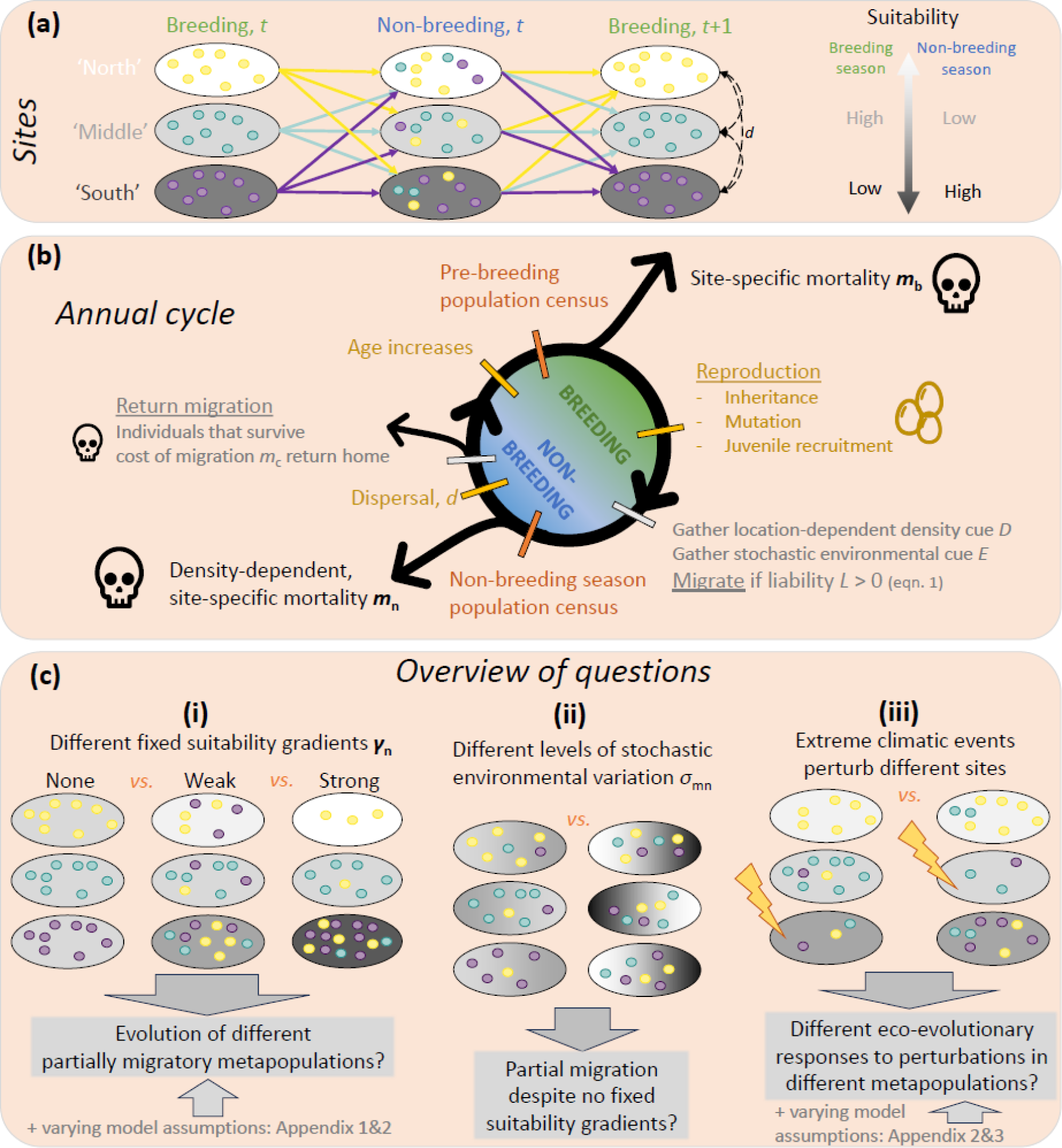
(a) Illustration of a partially-migratory metapopulation (PMMP) with S=3 sites (greyscale ovals) and subpopulations defined by individuals breeding in each site (small circles, color indicates breeding site). During the non-breeding season, individuals may stay in their breeding site or migrate to a different site (colored arrows), so that each site may contain a mixed group of residents and migrants from other sites. For the following breeding season, all individuals return to their home site, except a small fraction *d* of random dispersers (black dashed lines). Sites may differ according to a geographical breeding and/or non-breeding season suitability gradient. (b) Order of events in the annual cycle in our individual-based model. A year begins with all individuals increasing in age at the beginning of the breeding season (green), followed by the non-breeding season (blue). Events are color coded: Black: mortality events (with skulls). Grey: seasonal migration. Orange: population census. Gold: individual genes/properties change. Parameters in bold indicate vectors of site-specific values. (c) Overview of questions addressed. In (a) and (c), ovals with lighter *vs.* darker shades of grey represent sites less *vs*. more suitable during the non-breeding season (‘north’ *vs.* ‘south’).

Second, we should relax the assumption of a one-locus, two-allele genetic architecture. In many taxa, expression of migration versus residence is appropriately conceptualized as a quantitative genetic threshold trait (birds: Pulido 2011, fish: Dodson et al. 2013, mammals: Berg et al. 2019). Here, individuals express discrete alternative phenotypes when an underlying continuously distributed ‘liability’ falls below versus above a threshold (Roff 1996). Liability is assumed to depend on alleles at numerous loci of small effect (i.e. ‘infinitesimal model’), alongside various environmental effects. Substantial latent genetic variation may therefore exist, even among individuals expressing the same phenotype. This ‘cryptic’ variation can be revealed, and thus exposed to selection, when ECEs (or resulting reduced population densities) cause large environmental deviations in liabilities, such that new individuals cross the threshold and express the alternative phenotype (Pulido 2011, Reid and Acker 2022, Acker et al. 2023). Clearly, such quantitative genetic threshold traits will exhibit very different dynamics in the face of spatio-temporal environmental variation than simple Mendelian traits, or than traits that are continuously distributed on phenotypic scales. But, threshold trait architectures have not yet been fully built into eco-evolutionary theory.

Accordingly, we here present a general individual-based model designed to quantify eco-evolutionary dynamics of PMMPs in spatially structured seasonal landscapes, with migration formulated as a quantitative genetic threshold trait. First, we show that such PMMPs can readily maintain partial migration and underlying genetic variation, of forms and magnitudes that depend on spatial variation in non-breeding season environmental suitability. Second, we show that seasonal environmental stochasticity can promote partial migration, even in the absence of density-dependent constraints on vital rates. Third, we demonstrate how seasonal ECEs can induce strong eco-evolutionary dynamics, that emerge across diverging spatial and temporal scales. Our model and illustrative analyses therefore reveal key elements of how spatially structured and potentially mobile populations could respond to changing seasonal environments.

## Methods

Our model considers the full annual cycle of a (potentially) partially migratory metapopulation comprising subpopulations inhabiting *S* separate sites each with carrying capacity *K* individuals. Populations are sexually reproducing and age-structured with overlapping generations. The annual cycle comprises sequential breeding and non-breeding seasons, e.g. representing summer and winter (Fig. 1). Individuals can stay resident at their breeding site through the non-breeding season, or migrate to a different site, before returning home to their original site to breed the next year, or moving to a different site if dispersal occurs (with a small, fixed probability *d*). Local population densities and seasonal environmental characteristics (‘suitability’) at each site can affect vital rates during both seasons. Sets of sites can be constructed to caricature diverse landscapes. For example, we can envisage a northern temperate latitudinal gradient, with ‘northern’ sites providing good breeding season conditions and bad non-breeding season conditions and vice versa in the ‘south’, or other common patterns such as altitudinal (Spitz et al. 2018), coastal-to-inland (Allen et al. 2019), aridity (Serneels and Lambin 2001) or salinity gradients (Kasai et al. 2018), or ponds along a gradient of ephemeral to permanent (Grayson et al. 2011). Such environmental gradients can exist in either, neither, or both seasons.

Expression of migration versus residence is modelled as a threshold trait, where individual *i* migrates in year *t* if its liability *L_i_*(*t*) exceeds a threshold value *T*=0 (Fig. 1b). *L_i_*(*t*) comprises three components: a heritable additive genetic effect (i.e. ‘breeding value’) *a_i_*; a deterministic effect of local population density, *D*; and a stochastic effect of the environment, *E* (eqn 1). The value of *D* at site *j* at any time *t* is the ratio of the subpopulation size *n_j_*(*t*) to the carrying capacity *K_j_* (*D_j_*(*t*) = *n_j_*(*t*)/*K_j_*). The value of *E* is given by a random draw from a Gaussian distribution of mean 0 and variance 1, implemented at either the site level (“coarse-grained” environments, where all individuals in site *j* experience the same *E_i_* = *E_j_*), in our main results, or at the individual level (“fine-grained” environments, where each individual experiences a different *E_i_*), in Appendix S1. Liability-scale reaction norms are assumed to be linear, so for individual *i* in site *j* in year *t*,

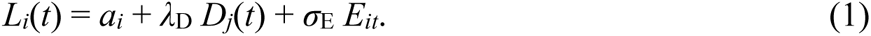

Here, the relative impacts of the non-genetic components *D* and *E* can be independently adjusted by the positive, non-evolving parameters *λ*_D_ and *σ*_E_, respectively set to 0.1 and 1 in our baseline scenarios, and both set to 0 in Appendix S1 such that liability is purely genetically determined. Note that values of *σ*_E_ and *λ*_D_ only determine interpretation of evolving breeding values relative to other scenarios, as liability-scale reaction norms are effectively dimensionless. Whereas *E* is a white-noise component and is equally often negative or positive, *D* is strictly positive and typically near 1 for a stable subpopulation near *K*.

Conditional on migrating, an unlinked, haploid destination gene determines where an individual goes. We assume that there is one allele that directs towards each site, and an individual’s allele can be that for any site other than its breeding site. Such large effect loci do exist for migratory tactics in nature (e.g. Sokolovskis et al. 2023). Since the destination gene is not expressed unless an individual migrates, selection on this locus is weaker the more resident a population is (Van Dyken and Wade 2010). If dispersal occurs, the destination gene mutates to ensure that, conditional on migrating, the individual moves to a different site from its new breeding site.

We apply adult mortality at three annual timepoints (Fig. 1b). First, a time-invariant but potentially site-dependent mortality probability *m*_b,*j*_ applies at the start of the breeding season. Second, a density-dependent mortality probability *m*_n,*j*_ applies at the start of the non-breeding season, following an exponential function of the number of individuals *n_j_*(*t*) currently present,

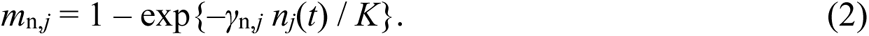

The strength of density dependence is modulated by the site-specific parameter *γ*_n,*j*_, representing local seasonal ‘suitability’. This mortality occurs independently of whether individuals that are currently present are resident at site *j* or migrated there from elsewhere. Note that we set *K* constant across sites and seasons, and implement seasonally and geographically varying site quality through varying ***m*_b_** and ***γ*_n_** (bold denotes a vector of site-specific values). Third, a constant mortality probability *m*_c_ applies at the end of the non-breeding season for individuals that migrated in the current year, hence representing a cost of migration. We let this cost be independent of the distance a migrating individual travelled, but in Appendix S2 we relax this assumption and weight the cost by the distance travelled (e.g., travelling between adjacent north and middle sites is less costly than travelling between more distant north and south sites).

Reproduction occurs after breeding season mortality. The number of offspring per individual depends on basal reproductive rate or ‘maximum fecundity’ *f* (expected number of recruited offspring per parent at low population densities, set to 3 in our main results, and 2 in Appendix S3), and the number of ‘slots’ available at the breeding site *j*, *K*–*n_j_*. Specifically, *n*_off_ offspring recruit, where *n*_off_=min(*K*–*n_j_*, *n_j_f*), such that post-breeding subpopulation sizes (juveniles + adults) cannot exceed *K*. We assume random and fully polygynandrous mating, sampling (with replacement) *n*_off_ mothers and *n*_off_ fathers among all surviving adults. Consequently, selfing is not prohibited, although the probability is tiny. All adults (age ≥ 1) are assumed capable of reproduction, although postponing age at maturity does not qualitatively alter our current conclusions (results not shown).

Inheritance and mutation of breeding values for liability to migrate is governed by the infinitesimal model (Barton et al. 2017), where an offspring’s breeding value *a_i_* is drawn from a normal distribution with mean equaling the mean of its parents’ breeding values, and variance equal to the initial subpopulation additive genetic variance, *V*_0_ (ignoring inbreeding, which has a negligible effect). Each offspring also randomly inherits one parent’s destination allele, such that the assumption of haploidy essentially captures a diploid system with no dominance. The locus mutates with probability *μ=*0.01, to one of the *S*–2 other possible alleles.

### Simulations

Our current eco-evolutionary simulations focus on systems with *S*=3 sites, generating the simplest possible PMMP where an individual can either stay resident through the non-breeding season or migrate to >1 alternative sites, although we additionally explore systems with *S*=5 sites (Appendix S2). Simulations are initiated with all subpopulation sizes at *K*, Poisson age distributions with mean 3, and Gaussian breeding value distributions with mean 0 (equaling the migration threshold) and variance *V*_0_=1. Each individual’s destination allele is randomly assigned, excluding the allele encoding each individual’s home site. Allele frequencies for each of the *S*–1 possible destinations are therefore approximately 1/(*S*–1) in each subpopulation.

We first quantified effects of different geographical gradients of seasonal suitability on the evolution and maintenance of PMMPs (Fig. 1ci). We independently set whether breeding season mortalities ***m*_b_** and/or non-breeding season suitabilities ***γ*_n_** exhibited weak or strong ‘north-south’ gradients, or else were equal across sites. We censused all individuals present at each site at two annual time points (Fig. 1b, red ticks), thus quantifying emerging spatio-seasonal population dynamics and underlying degrees and forms of migration. We further tested how PMMPs are affected by absence vs. presence of non-genetic effects on liability (*D* and *E* terms in eqn. 1), by fine- vs. coarse-grained environmental variation, and by distance-dependent vs. distance-independent costs of migration *m*_c_ (Appendix S1 & S2).

Second, we tested the ability of environmental stochasticity to generate partial migration in the absence of any deterministic geographical gradient in seasonal suitabilities (all ***γ*_n_**=1/3), by including random spatiotemporal variation in breeding and non-breeding season mortality (Fig 1cii). Specifically, for each site *j* in each year, a deviation drawn from a Gaussian distribution of mean 0 and standard deviation of *σ_m,_*_b_ or *σ_m,_*_n_, respectively, is added to the mortality probability (which we then bound between 0 and 1). Seasonal suitability is thus unpredictable and uncorrelated among sites and years. We also tested whether such environmental stochasticity could maintain partial migration even in the absence of any non-breeding season density-dependence (***γ*_n_**=0), a process often considered central to the evolution of stable partial migration.

Third, to quantify eco-evolutionary responses of PMMPs to specific extreme climatic events (ECEs), we imposed large perturbations on single sites (Fig. 1ciii). These one-off ECEs occur during the non-breeding season and kill 80% of all individuals present, independent of whether they were residents or had migrated there from elsewhere. We perturbed each site separately in PMMPs that had evolved under strong, weak or no suitability gradients (***γ*_n_**) and no stochasticity in non-breeding season survival (*σ*_m,n_=0). Such perturbations fall within the extremes observed in nature (e.g. Alonso-Andicoberry et al. 2002, Frederiksen et al. 2008). These simulations were chosen to clearly reveal general eco-evolutionary responses, as weaker ECEs (<80% mortality) produce weaker patterns but the same underlying mechanisms. We examined eco-evolutionary responses for small and large PMMPs (*S*=3 or 5), low and high intrinsic growth rate (*f*=2 or 3), and with or without distance-dependent costs of migration (Appendix S2 & S3).

Simulations were typically run for 10,000 years with 20 replicates per parameter set, to ensure that eco-evolutionary trajectories stabilize and to adequately capture among-replicate variation in outcomes. Simulations with stochastic mortality were run for 50,000 years, since evolutionary responses to such stochasticity can be slower and more subtle (Simons 2002). Seasonal population sizes were recorded yearly at each site (Fig. 1b), with non-breeding season counts representing the number of individuals currently present at the site (i.e., not the number of individuals breeding at a site that are currently alive). Individual-level characteristics (genotype, breeding site, migration destination, liability) were recorded every 50 years. ECEs occurred in year 9950, with individual-level data recorded every year for 50 years before and after, allowing detailed analysis of transient ECE-induced population and evolutionary dynamics (Appendix S1). All simulations were run in R version 4.2.1 (R Core Team 2022). Code is available online (Haaland 2023). All parameter values and definitions are summarized in Appendix S1: Table S1.

## Results

### Geographical gradients of seasonal suitability

Our first results demonstrate the general point that, as predicted given a multi-site structure with sufficiently large fixed differences among sites in non-breeding season suitability (i.e. gradients in strengths of density-dependence ***γ*_n_**), coupled with a quantitative genetic threshold trait architecture, variation in seasonal migration can readily evolve and be maintained (Fig. 2). Here, subpopulations breeding in sites that are more or less suitable in the non-breeding season became respectively less or more migratory, due to local adaptation in breeding values for liability to migrate (Fig. 2b,c). With no suitability gradient, fully resident populations evolve (Fig. 2a). As expected under the infinitesimal model, breeding values were in all cases normally distributed within subpopulations (Fig. 2d-f), and additive genetic variance stabilized to 2*V*_0_. Fully migratory, partially migratory and fully resident subpopulations are respectively characterized by breeding value distributions that evolved to be entirely positive, spanning zero, or entirely negative, achieved through evolutionary change in the means (Fig. 2d-f).

**Figure 2:**
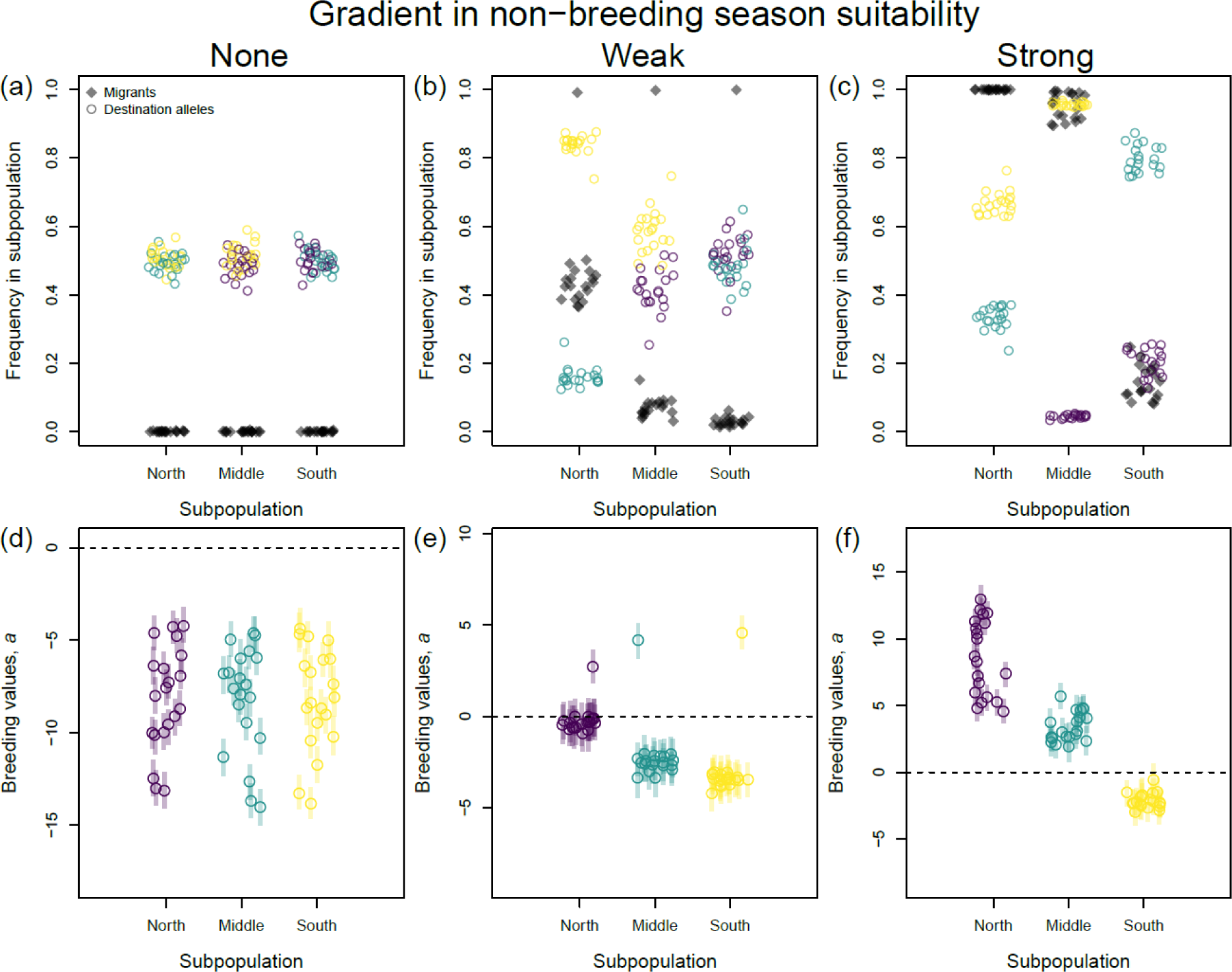
Metapopulation genetic structure after 10,000 years of evolution where sites experience (a, d) no differences in non-breeding season suitability, ***γ*_n_** = {1/3, 1/3, 1/3}, (b, e) weak differences, ***γ*_n_** = {1/2, 1/3, 1/5}, or (c, f) strong differences, ***γ*_n_** = {5, 1/2, 1/10}. (a-c): Migrant frequency (black diamonds) and allele frequencies at the destination locus (colored circles; purple, green and yellow respectively represent alleles coding for migrating to the north, middle and south site) in each subpopulation, averaged across 50 year intervals through the last 1000 years within each simulation. (d-f): Median (dark colored points) and interquartile range (light colored bars) of breeding values for migration liability in each subpopulation in year 10,000 for each replicate simulation. Dashed horizontal lines show the threshold liability *L*=0, above which an individual migrates; note differences in y-axis scales. All parameters except ***γ*_n_** set to baseline values (Appendix S1: Table S1).

Depending on costs of migration *m*_c_ and the steepness of the non-breeding season suitability gradient (***γ*_n_**), subpopulations can be partially migratory in the north and fully resident in the south (weak gradients, Fig. 2b,e) or fully migratory in the north and partially migratory in the south (strong gradients, Fig. 2c,f). This seemingly maladaptive migration in the south, and occasional fully migratory metapopulations (outliers in Fig. 2b,e), are caused by gene flow due to dispersal among subpopulations constraining local adaptation (confirmed by additional simulations with higher dispersal rates; results not shown).

With weak non-breeding season suitability gradients, PMMPs evolved migration rates and destinations such that non-breeding season geographical distributions of individuals generated similar density-dependent seasonal survival probabilities (Fig. 3b). Survival was slightly higher in the south site at equilibrium, but this site receives most seasonal migrants who also pay a cost of migration *m*_c_, contributing to evening out year-round survival probabilities. Thus, approximately balanced average fitness effects of residence and migration here emerge as an evolutionary endpoint, rather than being specified upfront with fixed parameter values. Non-breeding season survival probabilities differed more under strong suitability gradients (Fig. 3c), again possibly due to dispersal constraining local adaptation. In contrast, differences among sites in breeding season mortality (***m*_b_**) do not cause any adaptive responses. Simulations with suitability gradients in both seasons therefore generate PMMPs similar to those evolving with non-breeding season suitability gradients only (results not shown). Further, simulations with suitability gradients only in breeding season mortality generate fully resident metapopulations, mirroring the case with no suitability gradients (Figs. 2a,d & 3a,d).

**Figure 3:**
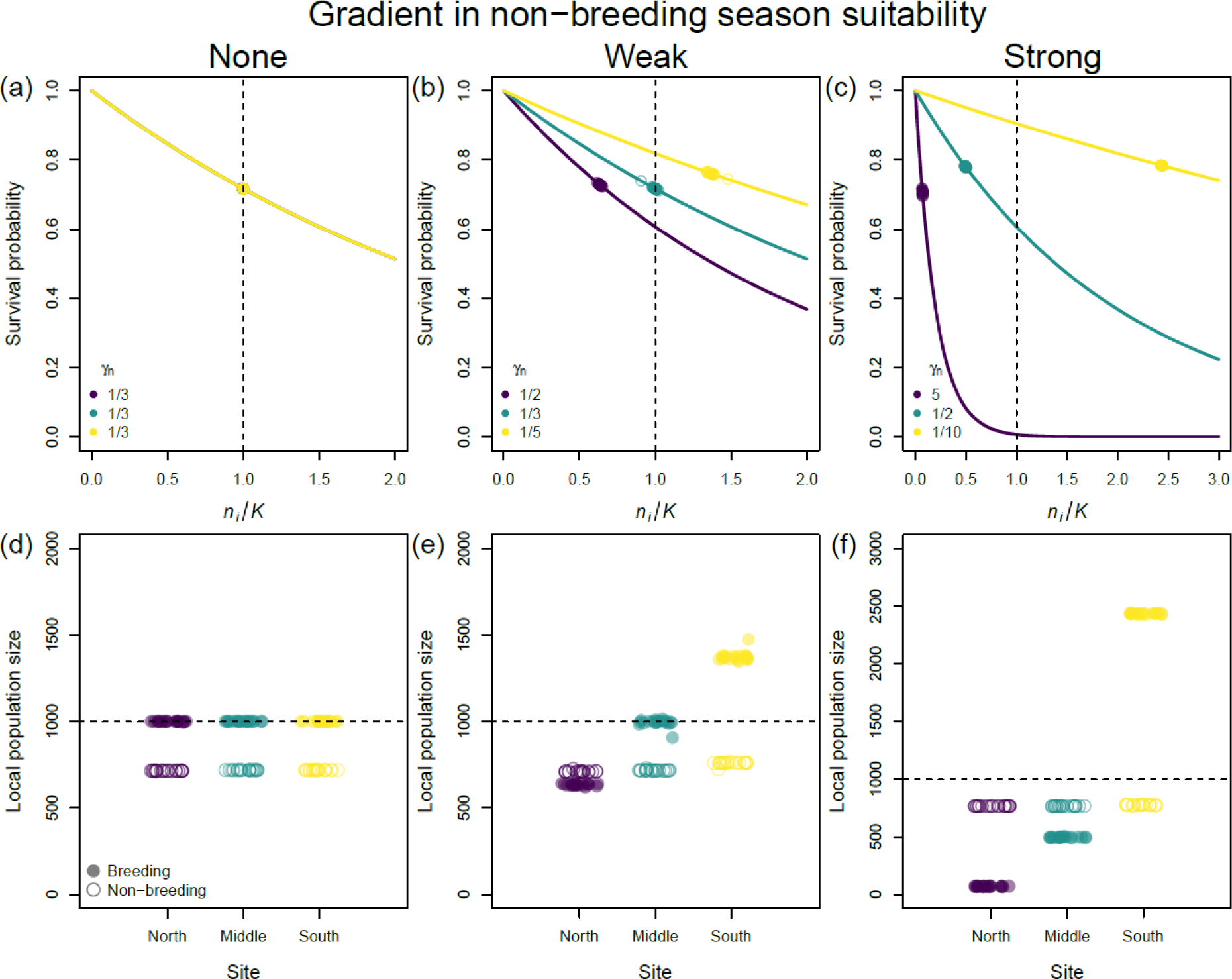
Metapopulation space use after 10,000 years of evolution where sites experience (a, d) no differences in non-breeding season suitability, ***γ*_n_** = {1/3, 1/3, 1/3}, (b, e) weak differences, ***γ*_n_** = {1/2, 1/3, 1/5}, or (c, f) strong differences, ***γ*_n_** = {5, 1/2, 1/10}. (a)-(c): Density-dependent non-breeding season survival (colored lines), i.e. the inverse of eqn. 2. Open circles along the lines show actual winter space use and survival probabilities averaged across the last 1000 years within each simulation; closed circles are means across 20 replicate simulations. With no non-breeding season suitability gradient (a), all lines and circles overlap, with yellow plotted last. Panels d-f show local population sizes during the breeding (filled circles) and non-breeding season (open circles) at each site, averaged across the last 1000 years within each simulation. Dashed lines show carrying capacity *K*. Note that y-axis scales differ. All parameters besides ***γ*_n_** set to baseline values (Appendix S1: Table S1).

The result that partial migration readily arises when sites differ in non-breeding season suitability remained robust when other model components were varied (Appendix S1), including whether the non-genetic components *D* and *E* (respectively local population density and stochastic environmental effects) contributed to migration liability or not, and whether environmental stochasticity *E* was coarse- or fine-grained. However, the presence and grain of variation in *E* increases among-year fluctuations in numbers of migrants versus residents, so that balanced fitness effects across strategies are only evident when looking across multiple years.

Qualitatively similar patterns also emerged given larger PMMPs (*S*=5 sites) and when including an effect of distance travelled on the mortality cost of migration (Appendix S2). While such a distance effect does not limit the conditions where partial migration evolves, it shifts the pattern of migration from ‘leapfrog’ to ‘chain’ (Lundberg and Alerstam 1986). Specifically, rather than migrants from northern subpopulations travelling all the way south, some northern migrants travel to middle sites, and higher non-breeding season population densities in those sites in turn cause an increased proportion of individuals from middle sites to travel south.

### Stochasticity in survival

Adding intermediate levels of stochasticity in non-breeding season survival can cause partial migration, even without any fixed spatial suitability gradient (Fig. 4a). Meanwhile, low stochasticity leads to full residence, while very high stochasticity can drive metapopulations extinct (Fig. 4a). Extinct metapopulations exhibited similar pre-extinction migratory frequencies as surviving metapopulations.

**Figure 4:**
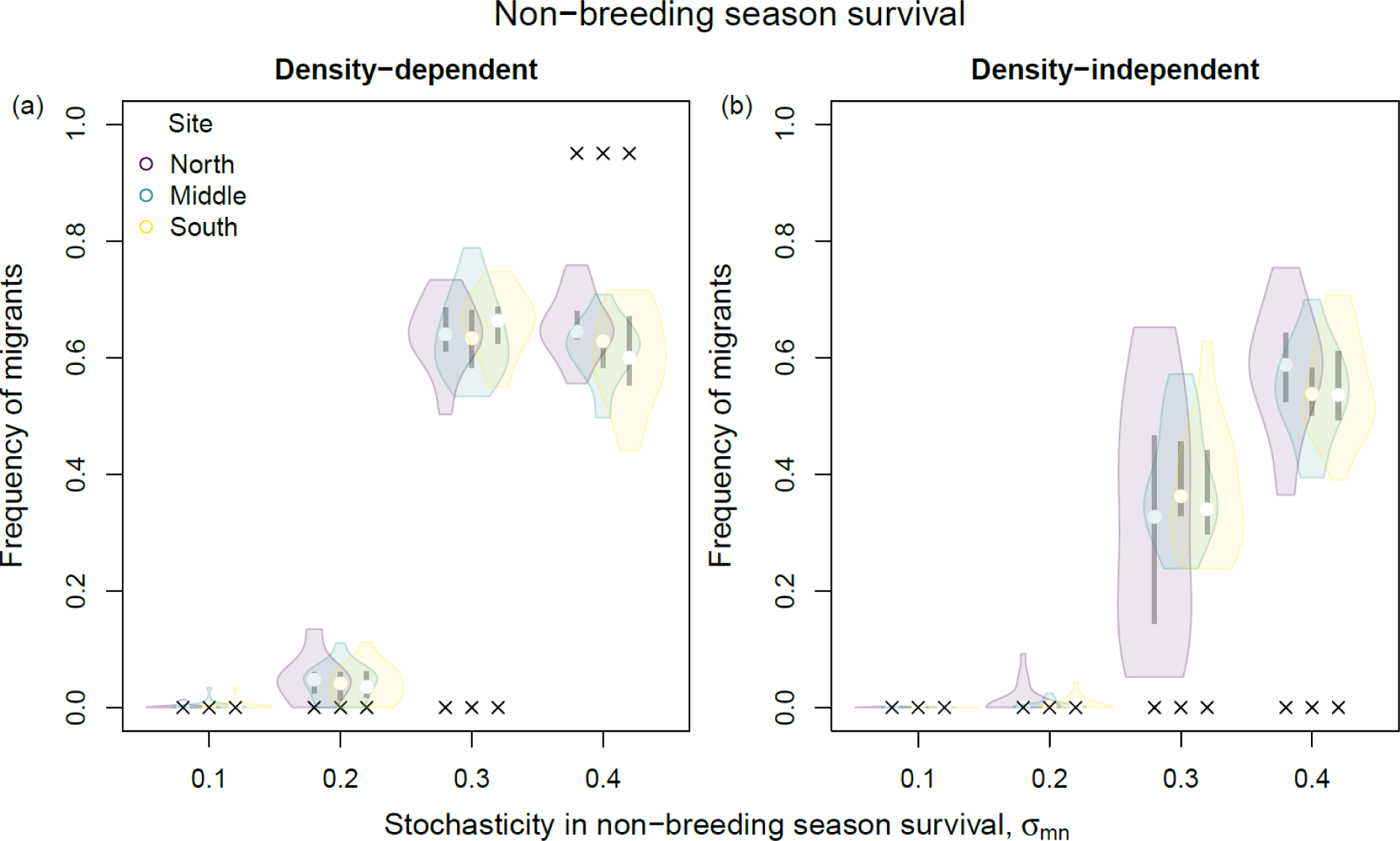
Evolved metapopulations with no fixed differences among sites in non-breeding season suitability, but stochastic variation in survival *σ*_mn_ among sites and years. (a) Non-breeding season survival is density-dependent, ***γ*_n_** = {1/3, 1/3, 1/3}. (b) Non-breeding season is density-independent, ***γ*_n_** = {0, 0, 0}. Shaded areas denote densities of the frequency of migrants at the end of 50,000 years of evolution for 20 replicate simulations, averaged across 50 year intervals through the last 1000 years, with colors representing subpopulation. Open circles represent medians, black boxes the interquartile range and black crosses the proportion of replicate simulations ending with subpopulation extinction. Plotted frequencies of migrants for extinct subpopulations (for *σ*_mn_=0.4 in panel a) are calculated across 50 year intervals through the last 1000 years prior to subpopulation extinction.

Further, beyond driving partial migration in the absence of among-site variation in non-breeding season suitability, environmental stochasticity can even generate partial migration in the absence of any density dependence at all (Fig. 4b). However, such PMMPs persisted only in narrow parameter space, as extinction occurred when stochasticity in survival was too high.

### Effects of extreme climatic events (ECEs)

Eco-evolutionary responses of PMMPs to non-breeding season ECEs depended on interactions between the location of the impacted site within the PMMP and the previously evolved migratory dynamics (Fig. 5; Appendix S2 & S3). Before the ECE (vertical dotted lines in Fig. 5) all subpopulations are stable below *K* (pre-breeding censuses are plotted, so any adult mortality through the annual cycle reduces population sizes). As is intuitive, non-breeding season ECEs impact the local subpopulation, and the whole PMMP, more strongly when more individuals are present at the impacted site (Fig. 5).

**Figure 5:**
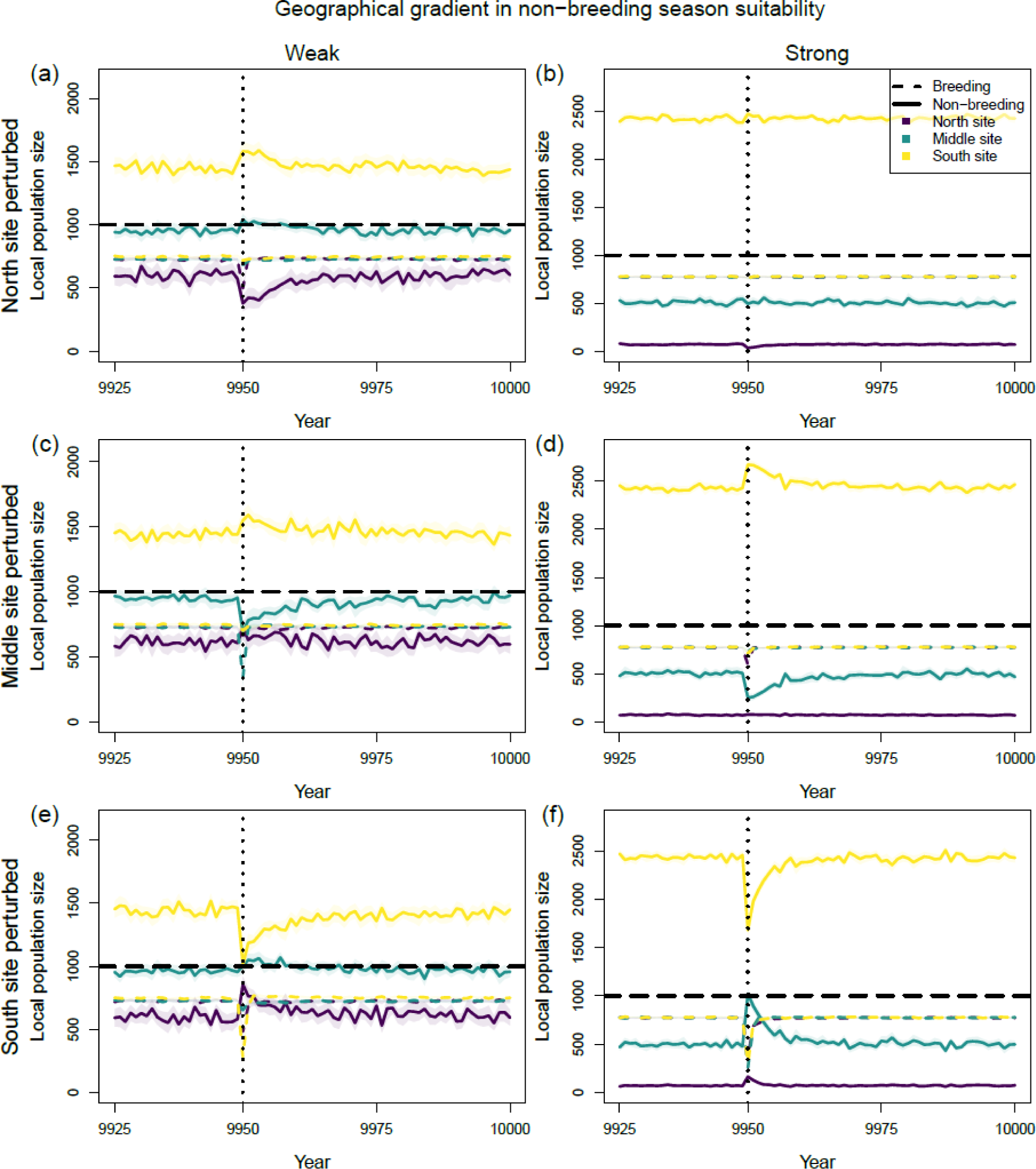
Population dynamics of evolved partially migratory metapopulations of 3 sites before and after an extreme climatic event (ECE) during the non-breeding season of year 9950 (vertical dotted lines), where the perturbation affects the (a, b) north, (c, d) middle, or (e, f) south site, given (a, c, e) weak (***γ*_n_** = {1/2, 1/3, 1/5}) or (b, d, f) strong (***γ*_n_**= {5, 1/2, 1/10}) non-breeding season suitability gradients. Breeding (dashed colored lines) and non-breeding season (solid colored lines) local population sizes are shown for each site. Thick colored lines and opaque bands represent means and 95% CI across 20 replicate simulations. Dashed black lines show carrying capacity. Other parameters set to baseline values (Appendix S1: Table S1).

For example, under strong non-breeding season suitability gradients, the north subpopulation is often fully migratory, and the north site receives no incoming migrants from other subpopulations, meaning that a local non-breeding season ECE has little effect on any subpopulations (Fig. 5b). In contrast, an ECE at the middle or southern site affects not only the local subpopulation by impacting local residents, but also the northern subpopulation(s) by impacting incoming migrants. The local subpopulation experiences an immediate decrease in both non-breeding and breeding season population sizes (Fig. 5c-f). The breeding population quickly recovers demographically, due to increased recruitment of juveniles as breeding season density dependence is relaxed. However, non-breeding season space use involves both residents and migrants, and needs to recover evolutionarily, which takes longer (about 20 years in the illustrated case, Fig. 5c-f). This is because the ECE causes three simultaneous evolutionary effects (illustrated in Fig. 6 for an ECE at the southern site).

**Figure 6:**
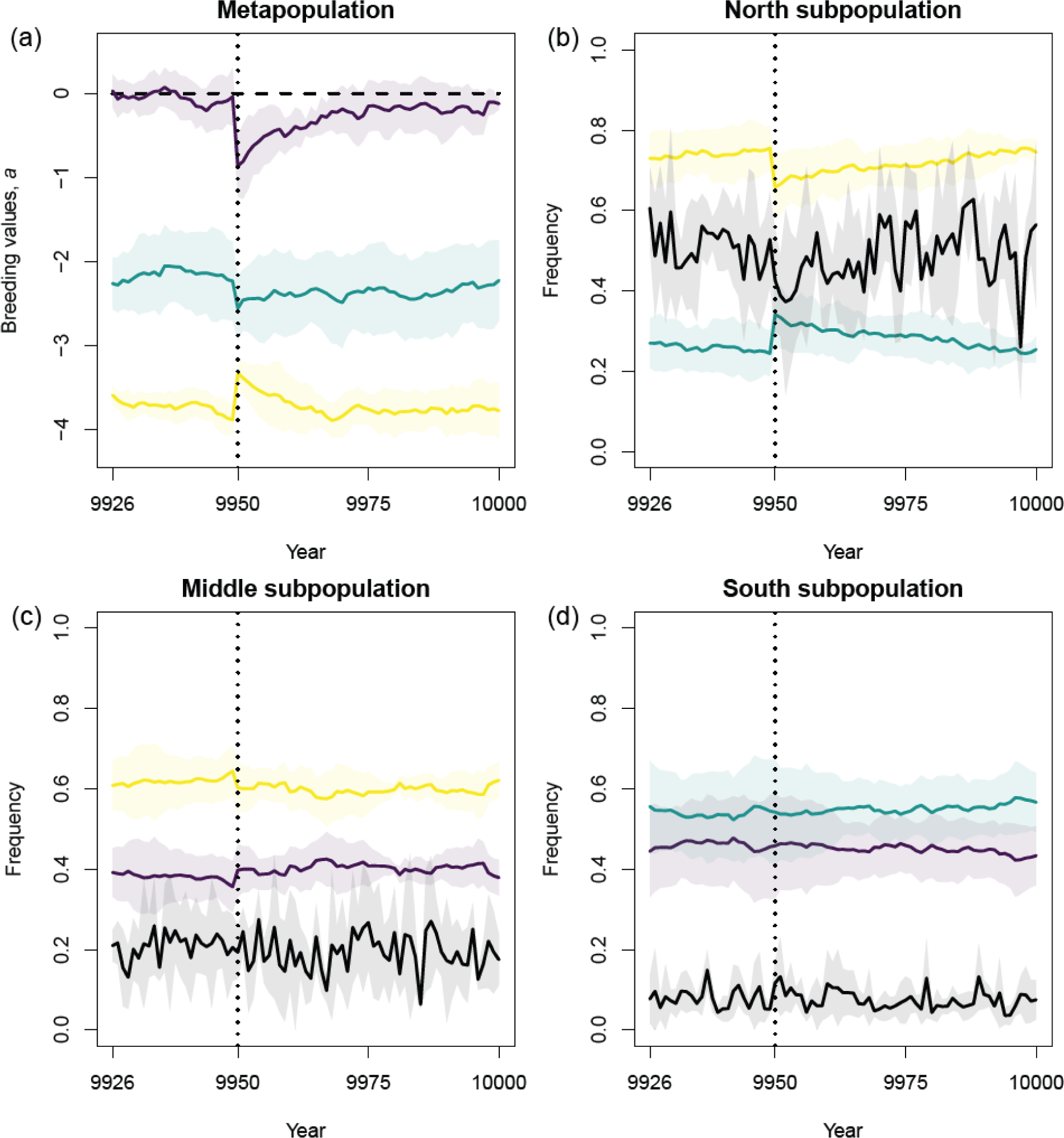
Evolutionary consequences of an extreme climatic event (ECE) striking the south site during the non-breeding season in year 9950 (vertical dotted lines), given a weak non-breeding season suitability gradient, ***γ*_n_** = {1/2, 1/3, 1/5} (as in Fig. 5e). (a) Subpopulation mean breeding across replicate simulations (opaque bands show 95% CI) before and after the ECE. Purple, green and yellow show respectively north, middle and south subpopulation. (b)-(d): For each subpopulation (panel header), colored lines show allele frequencies at destination locus (purple, green and yellow show respectively north, middle and south destination allele), and black lines migrant frequencies. Lines show means and opaque bands 95% CI across replicate simulations.

First, it causes strong selection against residence in the subpopulation breeding at the impacted site, increasing mean breeding values (Fig. 6a, yellow line). Second, it causes selection against migration in all other subpopulations, decreasing mean breeding values (Fig. 6a, purple and green lines). This selection is stronger in subpopulations from which more individuals migrated to the impacted site. Third, it causes selection against migrating to the impacted site, thereby altering frequencies of the destination alleles in all other subpopulations (Fig. 6b,c). The latter two effects are manifested as a sharp increase then gradual decrease in non-breeding season density in other sites following the ECE impact (Fig. 5d,f), due to higher proportions of local residents plus influxes of migrants from other subpopulations.

Together, these effects cause a medium-term decrease in non-breeding season subpopulation size in the perturbed site, which is balanced by concurrent increases elsewhere. Thus, while breeding season subpopulation sizes recover relatively fast (∼5 years), and can even temporarily overshoot pre-ECE densities (due to weaker non-breeding season density-dependence when fewer migrants arrive in the years following the ECE), non-breeding season space use takes much longer to return to the pre-ECE equilibrium (∼15-25 years, Fig. 5c-f).

The point that local ECEs can induce spatio-seasonal eco-evolutionary dynamics that extend far beyond the directly impacted site is further emphasized by considering larger PMMPs (*S*=5, Appendix S2). Here, an ECE that impacts the southernmost site, which receives many immigrants from all subpopulations given a strong non-breeding season suitability gradient, causes analogous strong population dynamic and evolutionary responses as observed above in all subpopulations. If the cost of migration *m*_c_ depends on distance travelled, more individuals from the northern subpopulations migrate to middle non-breeding season sites rather than the southernmost site. An ECE impacting the southernmost site then has slightly less impact on the breeding population size in the northernmost site than in our baseline scenario, but ripple effects in both population and evolutionary dynamics throughout the entire PMMP still persist.

The differing timeframes of ecological and evolutionary responses following ECEs, manifested as different times for breeding and non-breeding season population densities to return to pre-ECE levels, also remaine with lower fecundity (*f*=2 rather than 3) and hence reduced intrinsic population growth rate (Appendix S3). Alongside increasing population recovery times, reduced reproduction also slightly reduces the rate of evolution, thereby maintaining the discrepancy in recovery times for breeding versus non-breeding season population sizes.

## Discussion

The overarching ambition to predict spatio-temporal population dynamic responses to rapid environmental changes requires fully understanding the eco-evolutionary impacts of spatio-seasonal environmental variation on all vital rates. In partially migratory metapopulations (PMMPs), migration probability constitutes a vital rate, causing within-population variation in space use and responses to spatio-seasonal environmental variation (Reid et al. 2018, Payo-Payo et al. 2022). Our eco-evolutionary model, capturing multiple subpopulations that can be linked by migration formulated as a quantitative genetic threshold trait, demonstrates how both predictable and stochastic spatial environmental variation can readily generate stable partial migration. We also demonstrate how local seasonal shocks, for example caused by extreme climatic events, generate spatio-seasonal eco-evolutionary dynamics that percolate throughout the PMMP in time and space. Our model, and resulting insights on (meta)population responses to environmental perturbations, provide major advances towards building predictive, fully eco-evolutionary models of seasonally mobile natural populations.

### Evolution of stable PMMPs

PMMPs readily evolved in response to among-site differences in non-breeding season conditions (formulated as different strengths of density-dependence in mortality). Evolved metapopulation-level space use meant that individuals can experience similar survival probabilities despite occupying sites of different density and suitability, approaching ‘ideal free’ migratory patterns, as reported in natural PMMPs (e.g. Japanese sea bass *Lateolabrax japonicus*, Kasai et al. 2018; elk *Cervus canadensis*, Martin et al. 2022). Our geographical suitability gradients also generated instances of fully resident subpopulations occurring in sympatry with incoming migrants from other sites during the non-breeding seasons ‘shared non-breeding partial migration’ (Griswold et al. 2010). Further, stochasticity in non-breeding season survival among sites and years can drive evolution of partial migration, even without any variation in mean conditions, by effectively spreading the risk of experiencing harsh local conditions (Fig. 4a). This bet-hedging mechanism also works without any density-dependence in survival, if non-breeding season conditions are sufficiently unpredictable (high *σ*_mn_) and uncorrelated among sites and years (Fig. 4b). First stated by Cohen (1967), the point that stochasticity alone could drive partial migration has been largely omitted in later work, which focuses on non-breeding season density-dependence as a key mechanism (Kaitala et al. 1993, Taylor and Norris 2007). This focus may reflect the critique of bet-hedging as incompatible with frequency-dependent ESS models of partial migration, since resident individuals can outcompete partially migrant bet-hedgers during benign conditions interspersing unpredictable harsh years (Lundberg 1987).

In contrast, spatial variation in breeding season conditions did not drive partial migration in our current model. This is partly because we do not currently let breeding dispersal evolve, meaning that individuals experiencing poor breeding season conditions cannot leave. Exploring co-evolution of seasonal migration and breeding dispersal would certainly be of theoretical interest (e.g. Kokko and Lundberg 2001). However, individuals in many well-studied PMMPs are highly reproductively philopatric, despite consistent differences among sites in breeding season conditions (e.g. European shags *Gulosus aristotelis*, Barlow et al. 2013, Eurasian oystercatchers *Haematopus ostralegus*, Allen et al. 2019). Instead, differences in breeding season survival may cause more subtle eco-evolutionary effects on seasonal migration. For example, in our model, higher breeding season mortality leads to lower expected lifespan, but is compensated by higher per capita reproductive success. This is because juvenile recruitment is bounded by *K*, and more ‘slots’ for offspring are available if fewer adults survive. Resulting local adaptation due to life-history differences may affect whether evolved migratory strategies reflect bet-hedging adaptations (such as in Fig. 4), if alternative strategies differ in their fitness variance in stochastic environments (Starrfelt and Kokko 2012).

### Reponses to extreme climatic events (ECEs)

Our simulations highlight how major demographic perturbations caused by a single local ECE can cause far-reaching eco-evolutionary effects that extend throughout PMMPs. While local breeding population sizes typically recovered relatively fast following ECEs, impacts on non-breeding season population sizes persisted over much longer timeframes, often several decades. These outcomes result from simultaneous strong ECE-induced selection against residence in the subpopulation breeding at the perturbed site and selection against migration from other populations, generating spatially disruptive selection. Resulting evolutionary changes caused long-lasting ‘ripple effects’, since further evolutionary changes in both liability to migrate and in destination allele frequencies are required for the PMMP to return to its pre-perturbation equilibrium non-breeding season distribution. Such protracted periods away from system equilibrium following ECEs may help explain the lack of balanced fitness effects in a meta-analysis of populations with alternative migratory tactics (Buchan et al. 2020).

Seasonal environmental perturbations causing up to 80% local mortality have been documented in several partially migratory systems, including European shags *Gulosus aristotelis* during extreme winter storms (Frederiksen et al. 2008; Acker et al. 2021), greater flamingoes *Phoenicopterus roseus* during extremely cold winters or local toxic cyanobacterial blooms (Alonso-Andicoberry et al. 2002, Sanz-Aguilar et al. 2012), and bighorn sheep *Ovis canadensis* during disease epidemics (George et al. 2008) – although transmissible diseases have other, epidemiological consequences for metapopulation dynamics beyond those covered here (Balstad et al. 2021). At least some responses that arise in our simulations following perturbation are also known to occur in natural PMMPs. For example, after extreme winter storms on Isle of May, Scotland, the local European shag population became detectably more migratory, since selection increased mean breeding values in underlying liabilities (Acker et al. 2023). While similar analyses have not yet been performed on other subpopulations from which Isle of May hosts overwinter migrants, our model indicates that these subpopulations may be experience selection against migration (Fig. 6).

We envisage PMMPs in which migrants from one or more subpopulations share non-breeding season sites with residents. Such scenarios are common in nature, for example European shags in the north-eastern UK (Payo-Payo et al. 2022; Acker, et al. 2021), pied avocets *Recurvirostra avosetta* in France (Chambon et al. 2018), coastal-to-inland migrations of Eurasian oystercatchers *Haemotapus ostralegus* in the Netherlands (Allen et al. 2019), latitudinal or altitudinal migrations of songbirds such as blackcap *Sylvia atricopilla* (Berthold et al. 1992), robin *Erithacus rubicula* (Biebach 1983, Adriaensen and Dhondt 1990) and European blackbird *Turdus merula* (Zúñiga et al. 2017), ungulates such as bighorn sheep *Ovis canadensis* (Spitz et al. 2018), fin whales *Balaenoptera physalus* (Geijer et al. 2016) and wildebeest *Connochaetes taurinus* (Serneels and Lambin 2001), and among islands in tiger sharks *Galeocerdo cuvier* (Papastamatiou et al. 2013). Given that eco-evolutionary responses to weaker ECEs than we consider (<80% mortality) will be mechanistically similar but with smaller effects, there is thus a strong empirical basis on which to postulate that the population and evolutionary dynamics we capture, including eco-evolutionary consequences of ECEs, could widely occur in nature.

### Implications and future directions

Our general model formulation allows capturing diverse PMMP structures and ensuing vulnerabilities, including different ECE frequencies, spatial scales and durations, all of which are currently changing worldwide (IPCC 2023). While our current parameterisations and simulations are hypothetical and designed to highlight general principles rather than capture any particular system, we formulated our models so that all parameters can in principle be explicitly estimated from empirical data. For example, recent developments in quantitative genetic analyses allow estimation of additive genetic variance in liability to migrate, and associated seasonal selection (Acker et al. 2021a, 2023). Parameterizing and analysing full eco-evolutionary models of natural PMMPs will therefore become increasingly tractable, especially as tracking technologies become coupled with life-history and genetic data.

Such quantitative genetic analyses also allow partitioning temporary and permanent environmental variances in liability to migrate, and resulting phenotypic gene-by-environment interactions (Acker et al. 2023). Interestingly, evidence from diverse partially migratory systems indicates that much variation in expression of migration is attributable to permanent early-life effects: adult phenotypes are substantially fixed, with repeatabilities reaching 70-90% (Gillis et al. 2008, Grist et al. 2014, Chambon et al. 2019, Arnekleiv et al. 2022, Martin et al. 2022). Such permanent individual effects could exacerbate the size and duration of ECE effects on PMMP composition and dynamics, especially in long-lived species. Specifically, episodes of strong selection that remove sets of resident or migrant individuals will cause persistent shifts in phenotype frequencies due to phenotypic inertia alongside microevolutionary change (Acker et al. 2023). Thus, our current simulations may underestimate the full magnitudes and durations of eco-evolutionary dynamics that could occur. Incorporating evolution of such developmental plasticity, under varieties of coarse- and/or fine-grained environmental variation, is therefore a natural next step towards a better predictive understanding of the consequences of climate change and ECEs on PMMPs, and towards explaining why high migratory repeatabilities observed in nature may evolve in the first place.

A key remaining challenge for empiricists aiming to predict PMMP dynamics under altered regimes of spatiotemporal environmental variation is to estimate strengths and shapes of density-dependence in seasonal vital rates (Thorson et al. 2015), a central parameter in our and previous partial migration models. Our analyses also reveal that since the size and grain of environmental variation in liability to migrate can strongly affect conclusions on fitness differences among residents and migrants (Appendix S1), long-term studies are crucial for accurate inference. Fluctuating density-dependent selection may therefore have biased the observation of typically imbalanced fitness effects among migratory tactics in Buchan et al.’s (2020) meta-analysis, of which many studies focused on shorter time-scales. Finally, our model and associated parameterisations could be extended to consider (dis)assortative mating with respect to seasonal migration versus residence, which can affect mean levels of migration, individual plasticity, and population responses to environmental change (de Zoeten and Pulido 2020, Reid and Acker 2022). Yet, while some assortative mating might be expected, at least some partially migratory systems show limited or no evidence of phenotype-dependent assortative mating (Grist et al. 2017, Méndez et al. 2020). Models with random mating may therefore provide reasonable approximations. Our current model and results thus provide important steps towards understanding how natural PMMPs may respond to current dramatically changing spatio-seasonal environmental conditions.

## Acknowledgements

We thank Sarah Wanless, Cassandra Ugland and Ellen Martin for useful discussions. This work was funded by Research Council of Norway grants 223257 and 313570.

## Author contributions

TRH and JMR conceived the objectives. TRH built and analysed the model with input from JMR, APP, PA and IIR. TRH wrote the manuscript with input from JMR. All authors contributed to conceptual development, manuscript editing and presentation.

## Appendix S1: Baseline model scenarios and variations

Here we summarize all default parameter values (Table S1) and the baseline model scenarios, then illustrate the effects of varying two central assumptions (Fig. S1 & S2).

### Summary of model scenarios

#### Landscape size

All our eco-evolutionary scenarios shown in the main text involve metapopulations spanning *S*=3 discrete sites. In Appendix S2 we explore scenarios with *S*=5 sites.

#### Liability components

As shown in eqn. 1, liability to migrate can consist of a genetic component *a*, a stochastic environmental component *E*, and a density component *D*. The size of the two non-genetic liability components can be independently varied by parameters *σ*_E_ and *λ*_D_, respectively. Our baseline assumption is that these components are both non-zero (default parameter values are set to *σ*_E_=1 and *λ*_D_=0.1), but below (Fig. S1a-c, S2b) we set them both to 0, letting liability be exclusively genetically determined. We also assume coarse-grained variation in the environmental components (i.e., that different random values are drawn only for each site) as a baseline, but below (Fig. S1d-f. S2c) we let this variation be fine-grained (i.e., each individual draws a different random value). Thus, Figure S1 & S2 are the only places where the baseline assumptions are altered.

#### Stochasticity in survival

As a baseline we assume no stochasticity in seasonal survival probabilities, except in main text section “Results: Stochasticity in survival” (Fig. 4) in which stochasticity in non-breeding season mortality (*σ*_m,n_) is varied. We do not show results for stochastic variation in breeding season mortality (*σ*_m,b_), since neither deterministic nor stochastic variation in breeding season mortality has evolutionary consequences in our model given that dispersal rates cannot evolve.

#### Distance-dependent costs of migration

Our baseline assumption is that costs of migration are independent of the distance travelled among seasonal locations, but we explore the consequences of making this mortality cost distance-dependent in Appendix S2 where *S*=5, as such an effect is more evident in larger landscapes. Both detailed analyses of evolved partially migratory metapopulations (PMMPs), and their responses to extreme climatic events (ECEs), are shown.

#### Maximum fecundity

Finally, we alter the population intrinsic rate of increase by changing the maximum fecundity *f* in Appendix S3, from 3 offspring (baseline assumption, everywhere else) to 2 (Appendix S3), examining any effects of reduced growth rate on population and evolutionary dynamics following ECEs.

**Table S1:** List of model parameters and notation used, alongside descriptions and default values or range explored. Bold parameter names indicate a vector of values.

| Parameter | Description | Default value or range |
| --- | --- | --- |
| $S$ | Number of sites | 3 (baseline) or 5 (Appendix S2) |
| $K$ | Carrying capacity (per site), number of individuals | 1000 |
| $n_j$ | Local population density in site $j$. Varies seasonally. During the non-breeding season, $n_j$ counts both residents from the local subpopulation (breeding in site $j$), and any incoming migrants from other subpopulations (breeding in sites $\neq j$) | Positive. Bounded by $K$ during the breeding season.<br>Unbounded during the non-breeding season |
| $L_i$ | Liability to migrate for individual $i$. A quantitative genetic threshold trait. Given by Eqn. 1 | Continuous, unbounded |
| $T$ | Threshold for seasonal migration – individual migrates if $L > 0$ and remains resident if $L < T$ | 0 |
| $a_i$ | Additive genetic component (‘breeding value’) of migration liability for individual $i$ | Continuous, unbounded |
| $D_j$ | Density-related component of migration liability in site $j$, $n_j/K$ | Continuous, positive |
| $E_i, E_j$ | Stochastic environmental component of migration liability, $E \sim \text{Normal}(0,1)$, drawn per individual $i$ (fine grain) or per site $j$ (coarse grain) | Continuous, unbounded |
| $\lambda_D$ | Scalar for the density component to autumn migration decision | 0.1 (baseline) or 0 (Appendix S1) |
| $\sigma_E$ | Scalar for the stochastic environmental component to autumn migration decision | 1 (baseline) or 0 (Appendix S1) |
| $\mu$ | Destination gene mutation rate | 0.01 |
| $V_0$ | Initial subpopulation additive genetic variance in $a$ | 1 |
| $f$ | Maximum fecundity | 3 (baseline) or 2 (Appendix S3) |
| $d$ | Dispersal rate | 0.01 |
| $\mathbf{m_b}$ | Breeding-season mortality, a vector of length $N$ | No gradient (default):<br>{0.1, 0.1, 0.1}<br>Gradient: {0.05, 0.1, 0.15} |
| $\mathbf{m_n}$ | Density-dependent non-breeding season mortality. Given by Eqn. 2. | [0, 1] |
| $\gamma_n$ | ‘Suitability’ – parameter affecting strength of density-dependence in non-breeding season mortality, a vector of length $N$ | No gradient: {1/3, 1/3, 1/3}<br>Weak gradient: {1/2, 1/3, 1/5}<br>Strong gradient: {5, 1/2, 1/10} |
| $\sigma_{mb}$ | Size of stochastic component to breeding season mortality (standard deviation) | 0 |
| $\sigma_{mn}$ | Size of stochastic component to non-breeding season mortality (standard deviation) | 0 (baseline), 0.1, 0.2, 0.3 or 0.4 (Fig. 4) |
| $m_c$ | Mortality cost of migration | 0.02 |
| $t_{\max}$ | Number of time steps simulations run for | 10 000 or 50 000 |

### Effects of non-genetic liability components and environmental grain

As Figure S1 shows, equilibrium metapopulation characteristics remain largely unchanged when varying our baseline assumptions about non-genetic liability components and grain of environmental variation. Setting *σ*_E_ and *λ*_D_ to 0, or applying fine-rather than coarse-grained environmental variation, both still produce local adaptation in breeding values for migration liability (Fig. S1a, b) and destination genes (Fig. S1c, d: colored points), with the same resulting proportions of migrants (Fig. S1c, d; black diamonds) and non-breeding season distributions (Fig. S1e, f; open circles) as in our baseline scenarios (main text and everywhere else).

**Figure S1.**
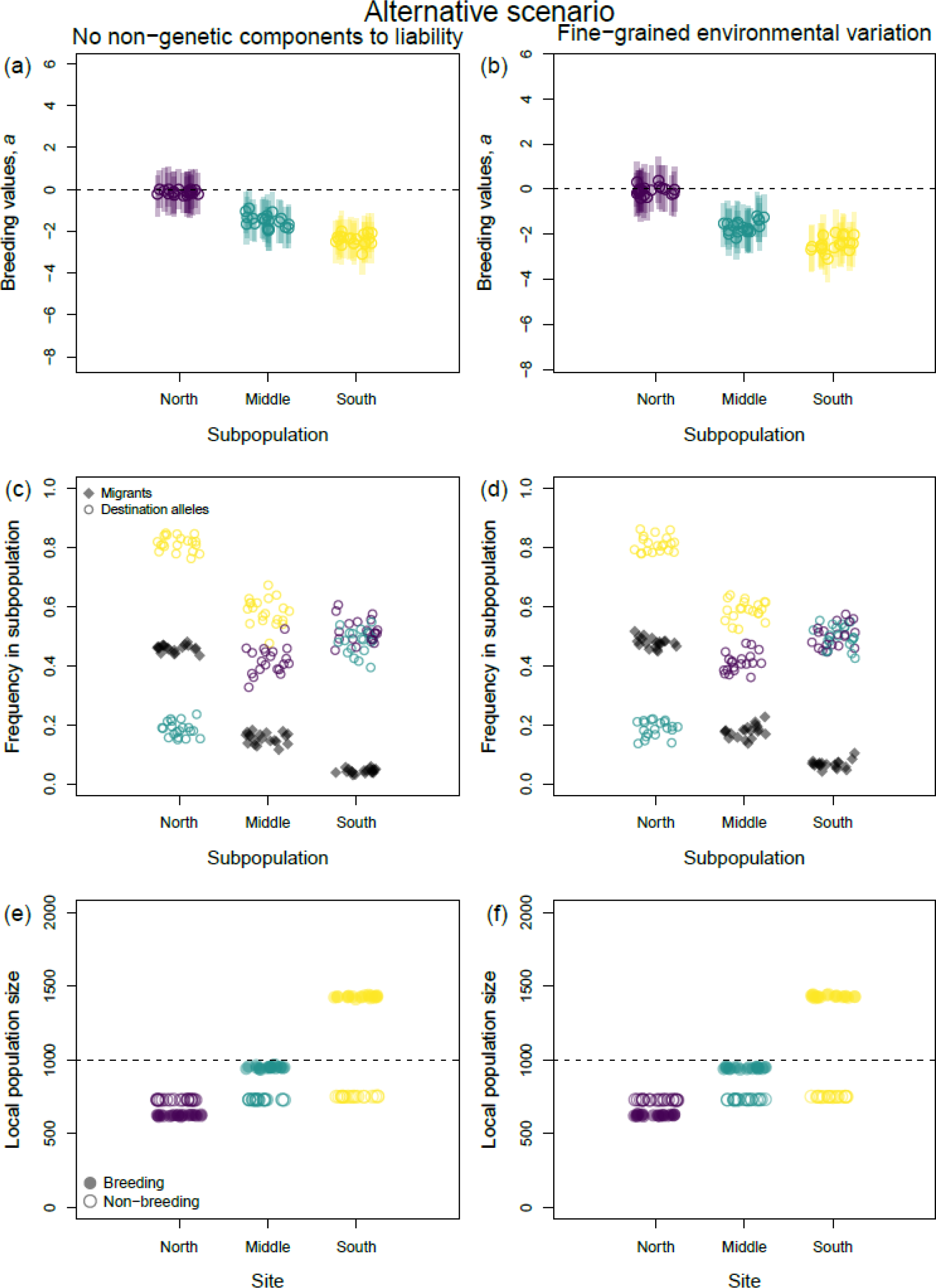
Metapopulation characteristics after 10,000 years of evolution under two sets of alternative model assumptions. (a, c, e): No non-genetic components affect migration liability, i.e., *λ*_D_=0 and *σ*_E_=0. (b, d. f): Fine-grained environmental variation, i.e. environmental components of migration liability vary at the individual level *E_i_* rather than the site level *E_j_*. In either case, all other parameters are set to baseline values (Appendix S1: Table S1) and landscapes exhibit a weak gradient of non-breeding season suitabilities, ***γ*_n_** = {1/2, 1/3, 1/5}, i.e. as in the middle columns of main text Figs. 2 and 3. (a, b): Median (dark colored circles) and interquartile range (light colored bars) of breeding values for migration liability in each subpopulation, for each replicate simulation. Dashed horizontal lines show the threshold liability *L*=0, above which an individual migrates. (c, d): Frequency of migrants (black diamonds) and destination alleles frequencies (colored circles; purple, green and yellow respectively represent alleles coding for migrating to the north, middle and south site) in each subpopulation. Frequencies are averaged across 50 year intervals through the last 1000 years within each simulation. (e, f): Local population sizes during the breeding (filled circles) and non-breeding season (open circles) at each site, averaged across the last 1000 years within each replicate simulation. Dashed horizontal lines show the carrying capacity *K*.

**Figure S2:**
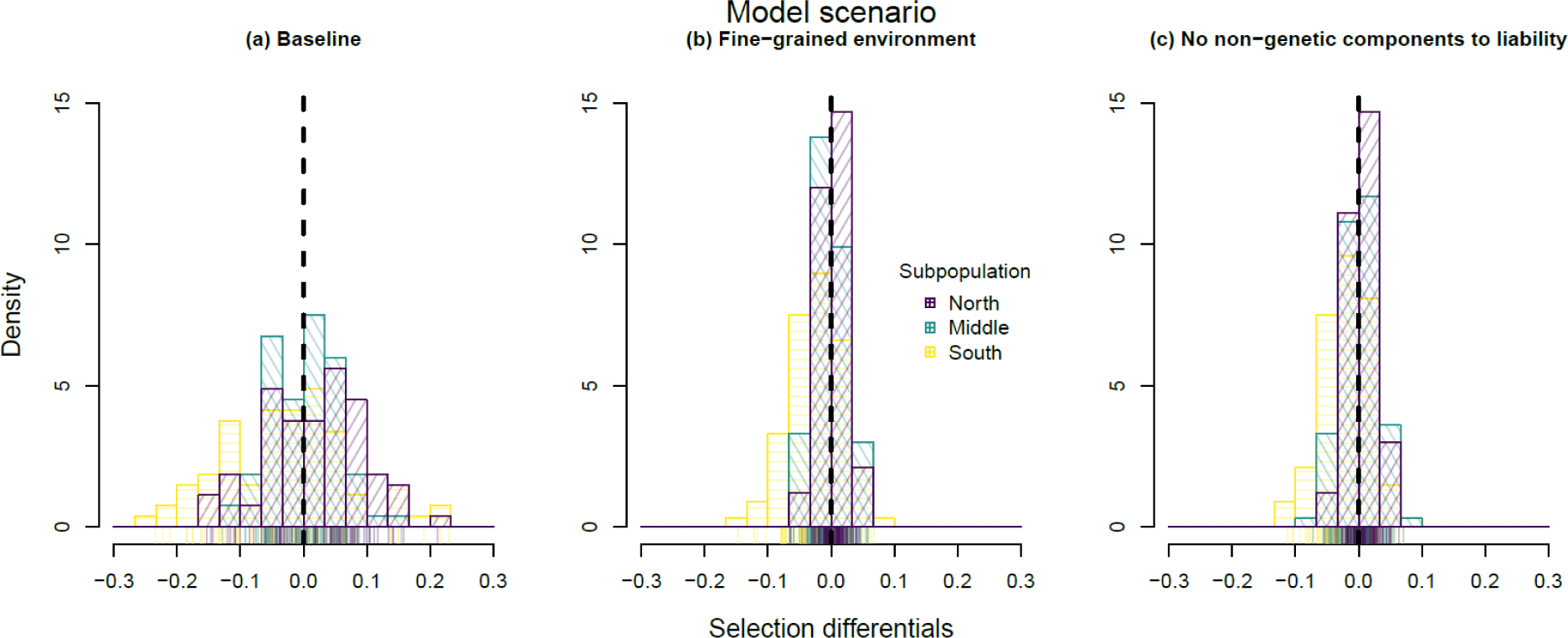
Histograms of yearly survival differentials on residence vs. migration per subpopulation (colors, fill hatching) at the end of 10,000 years of PMMP evolution. We consider weak geographical gradient in non-breeding season suitability (***γ*_n_** = {1/2, 1/3, 1/5}, i.e. as in the middle columns of main text Figs. 2 and 3), with three scenarios: (a) baseline parameters (see Appendix S1: Table S1), (b) fine-grained variation in environmental liability components, and (c) no non-genetic liability components. We obtain up to 100 semi-independent observations per subpopulation, calculating new selection differentials 20 times for 5 randomly chosen replicate simulations at 5 year intervals in the last 1000 years, unless the particular subpopulation is fully resident or fully migratory in a certain year. X-axis ticks show individual observations and histograms show densities (bin area summing to 1). Positive values of selection differentials (i.e. survival of residents minus survival of migrants) indicate that residents to better, negative values indicate that migrants do better. Dashed 0 line indicates equal survival among tactics.

However, differences among the scenarios do emerge regarding variation in migratory tactics exhibited among years. Specifically, scenarios with non-genetic components to liability (non-zero *σ*_E_ and *λ*_D_) produce subpopulations with much greater among-year variation in frequencies of migrants, especially for subpopulations with breeding value distributions centered around the threshold value *T*=0 (causing 50/50 residence/migration on average). These fluctuations also cause large year-to-year fluctuations in the fitness consequences of residence vs migration, since more or fewer individuals than expected may migrate in any given year, causing local densities and thus density-dependent survival rates in resident sites or migratory destinations to vary (wider distributions in Fig. S2a). This effect is especially pronounced for the subpopulation in the site with the most benign non-breeding season conditions (south subpopulation, yellow & horizontally hatched histogram in Fig. S2), where fitness benefits of residence vs migration vary greatly depending on how many visiting migrants there are from northern subpopulations. In contrast, fine-grained or no environmental variation in liabilities causes much smaller year-to-year fluctuations in migration rates and thus density-dependent survival (Fig. S2b, c), although fluctuations in selection are still greatest for the south subpopulations.

## Appendix S2: Movement and metapopulation evolution in larger landscapes

Here we show some cases of eco-evolutionary dynamics in larger partially migratory metapopulations (PMMPs), letting the metapopulation span *S*=5 discrete sites, rather than 3 as in the main text and elsewhere. We first illustrate how letting the cost of migration *m*_c_ depend on the distance travelled between sites can affect PMMP evolution (Fig. S3). Next, we show how the effects of extreme climatic events (ECEs) in larger PMMPs are analogous to those occurring in the smaller PMMPs shown in the main text, both without (Fig. S4) and with (Fig. S5) distance-dependent costs of migration.

### Effect of distance travelled on cost of migration

In order to implement an effect of the distance travelled between breeding and non-breeding season sites on the cost of migration (*m*_c_, a survival cost) whilst keeping overall mortality constant (*m*_c_=0.02), we let the random draw among all migrants (where a fraction *m*_c_ is to suffer mortality) be weighted by the distance they travelled between sites. We essentially assume that all sites are arranged in a straight line at equally spaced intervals of 1, such that e.g. an individual travelling from the north to the south patch when *S*=3 covers a distance of 2, whereas when *S*=5 it must cover a distance of 4. Costs of migration do not vary with ‘latitudinal’ position (i.e. travelling from north to middle sites is equally costly as travelling from south to middle sites).

Such a distance-dependent cost of migration causes subtle but noticeable shifts in migratory behaviour and spatio-seasonal PMMP structure relative to our baseline scenarios with distance-independent costs (Fig. S3, left (distance-independent cost) versus right (distance-dependent cost) column). With *S*=5 sites and a strong geographical gradient in non-breeding season mortality (Fig. S3c) all subpopulations except the southernmost become fully or partially migratory, with allele frequencies at the Destination locus strongly favoring migration to the southernmost site. As with *S*=3 sites, the southernmost subpopulation has fewer or no migrants, and thus experiences much less selection at the Destination locus – although there is a bias towards going to the middle-south site, which is also the second most favored destination for other subpopulations. Under weak gradients in non-breeding season mortality (Fig. S3a), all subpopulations are partially migratory but with much lower frequencies of migrants, with only the northernmost having more than 50 % migrants in most years. Selection at the Destination locus is therefore weaker, with only north subpopulation showing a clear preference for going to the south site. No preference in migratory destinations is evident in the middle, middle-south or south subpopulations.

When making costs of migration distance-dependent, the proportion of migrants from northern populations going all the way to the southernmost site is somewhat reduced. This causes (in Fig. S3 b, d) slightly more migrants from the two northern subpopulations to the middle sites, in turn increasing non-breeding season population densities in the middle sites, causing an increase in migrant frequencies in those subpopulations. Furthermore, whereas selection on the Destination locus is less strong in the middle subpopulations in panels a and c, this increasing migrant frequency now strengthens selection in these subpopulations, causing a clearer signal in panels b and d of the southernmost site being the favored migratory destination and the other sites being disfavored. Thus, incorporating an effect of the distance travelled to the cost of migration causes a slight shift of migratory pattern from “leapfrog migration” towards “chain migration”, as expected from previous theory (Lundberg and Alerstam 1986).

**Figure S3:**
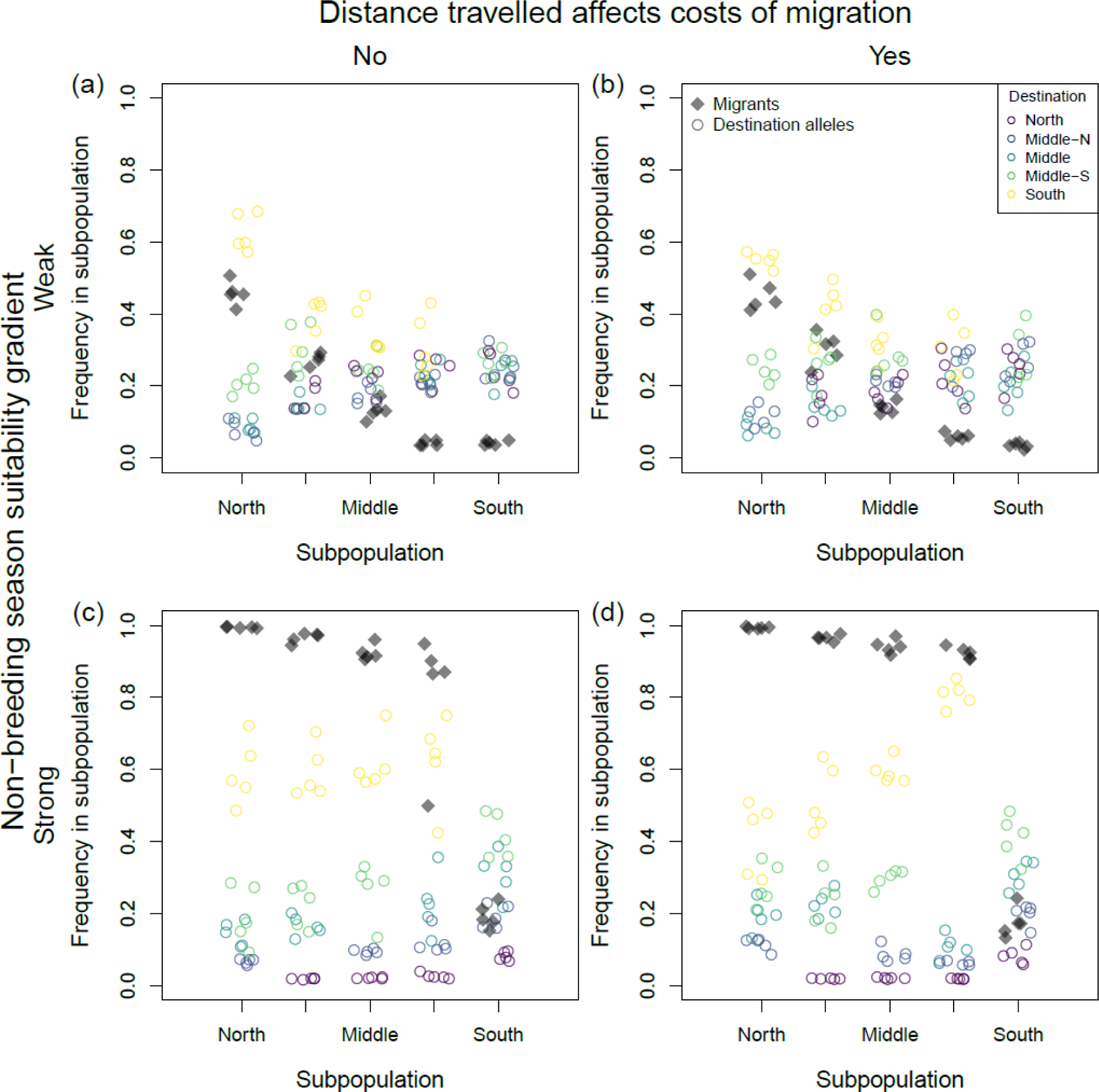
Metapopulation structure after 10,000 years of evolution where *S*=5 sites experience a weak (a, b) or strong (c, d) gradient in non-breeding season suitability, and where costs of migration do (b, d) or don’t (a, c) depend on the distance travelled among sties. Migrant frequency (black diamonds) and allele frequencies at the destination locus (colored circles) for each subpopulation are shown, averaged across 50 year intervals through the last 1000 years within each replicate simulation. The strong gradient of non-breeding season suitabilities uses values of parameter ***γ*_w_** = {5, 1, ½, 1/3, 1/10}, and the weak gradient uses ***γ*_w_** = {1/2, 2/5, 1/3, 1/4, 1/5}.

### ECEs in five-site PMMPs

Population and evolutionary dynamics of PMMPs with *S*=5 sites following non-breeding season ECEs follow similar principles as those occurring in smaller PMMPs (Fig. 5-6). As stated in the main text, a non-breeding season ECE causes three concurrent metapopulation-wide evolutionary responses, the strength of which all depend on the non-breeding season local density of individuals in the perturbed site (Fig. 6). These are:

i. Strong selection against residence and for migration occurs in the subpopulation breeding at the perturbed site,
ii. Selection against migration and for residence occurs in all other subpopulations, decreasing mean breeding values, more strongly so for subpopulations with more migrants overwintering in the perturbed site
iii. Selection on the destination locus in all other sites than the perturbed site against going to the perturbed site, again more strongly in subpopulations with more migrants overwintering in the perturbed site. In the subpopulation breeding at the perturbed site, selection on the Destination locus does not occur directly in response to the ECE, but indirect selection may occur in the years after the ECE due to altered overwintering densities in the other sites following changes in population densities and migratory strategies of other subpopulations in the PMMP.

For population dynamics, effects are also greatest if more individuals are present in the perturbed site during the non-breeding season. With a strong geographical gradient in non-breeding season suitability, the northernmost subpopulation is fully migratory, and thus an ECE at this site has no effect (Fig. S4b). But, with a weaker suitability gradient some individuals remain resident year round, and the perturbed site experiences a decrease in breeding population size, a decrease in non-breeding season population size, and southern sites experience an increase in non-breeding season population size (because of an increase in migratory liability, effect (i) above). If ECEs perturb sites that also receive incoming migrants from other sites (Fig. S4c-f), the abovementioned effect (i) occurs, along with effects (ii) and (iii) causing evolutionary shifts in the Destination locus as well. Selection against migrating to the perturbed site and for migrating to other sites is typically stronger in subpopulations the larger the proportion of migrants who would go to the perturbed site at the time of the ECE. Resulting non-breeding season PMMP space use is thus shifted following the ECE towards other less suitable sites, with this suboptimal metapopulation distribution lasting potentially for decades while the evolutionary effects (i-iii) return to pre-ECE equilibrium.

Comparing Figures S4 versus S5 reveals the population dynamics of five-site metapopulations in response to a non-breeding season ECE in PMMPs that have evolved without versus with an effect of distance travelled on the cost of migration. As shown in Fig. S3, including an effect of distance causes a shift from ‘leap-frog’ migration to ‘chain’ migration – i.e., migrants from more northern populations to migrate middle sites rather than the southernmost sites, in turn causing more individuals in the middle populations to migrate south. In the population dynamics time series, this can be observed for example in the d panels in Figs. S4 and S5: A non-breeding season ECE striking the middle site has little effect on northern populations in Fig. S5 (because northern migrants travel cost-free all the way south), but has a notable effect on northern breeding population sizes in Fig. S5 (downtick of darker dashed lines at year 9950).

**Figure S4:**
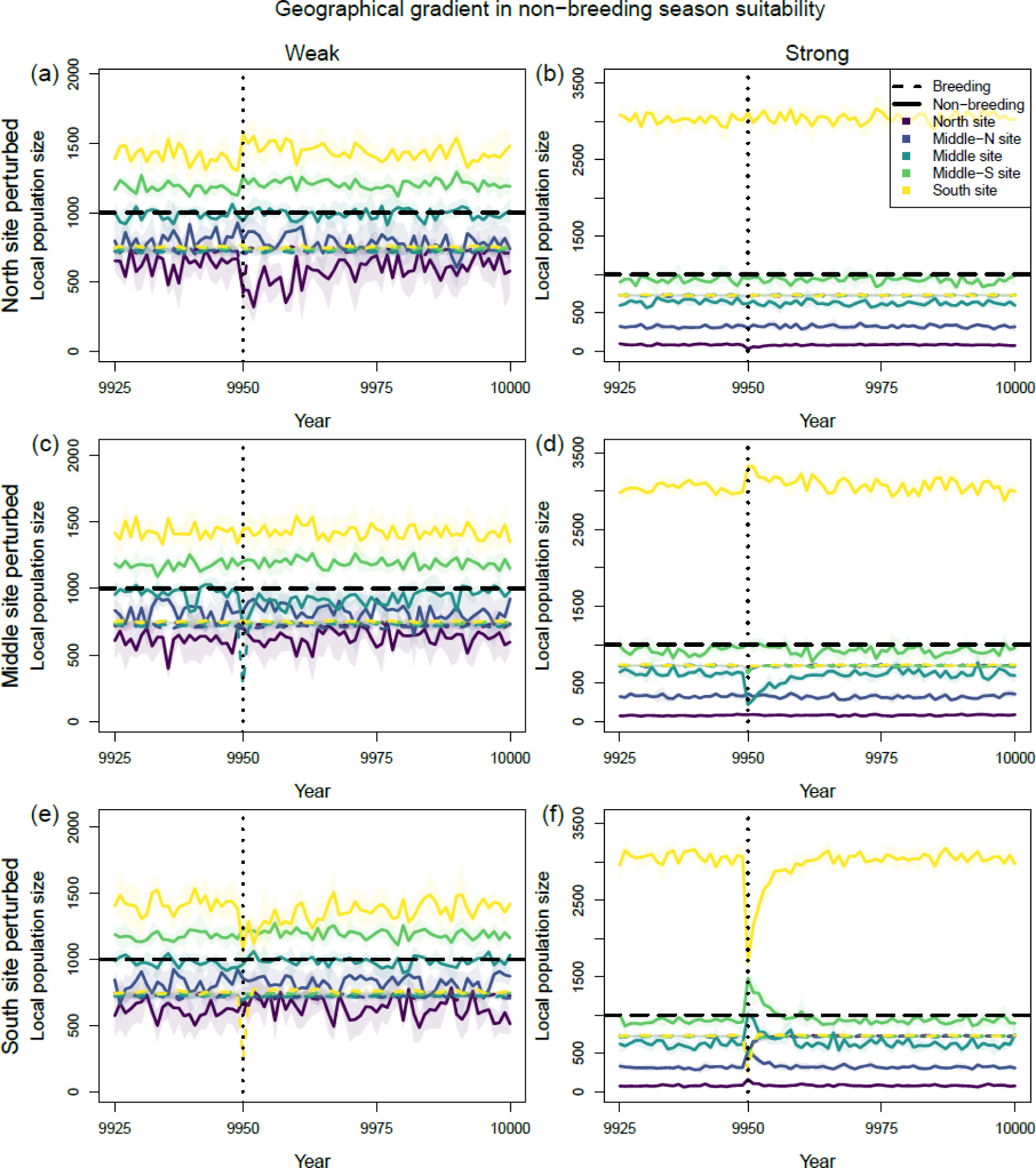
Population dynamics of evolved partially migratory metapopulations with 5 subpopulations before and after a non-breeding season ECE at year 9950 (vertical dotted lines). Breeding (dashed colored lines) and non-breeding season (solid colored lines) local population densities are shown for each site. Thick colored lines and opaque bands represent means and 95% CI across replicate simulations. Dashed black line represents carrying capacity. (a, c, e): Weak geographical gradient in non-breeding season suitability (density-dependent mortality parameter ***γ*_n_** = {1/2, 2/5, 1/3, 1/4 1/5}). (b, d, f): Strong geographical gradient in non-breeding season suitability (density-dependent mortality parameter ***γ*_n_** = {5, 1, 1/2, 1/3 1/10}). (a, b): Northernmost site perturbed. (c, d): Middle site perturbed. (e, f): Southernmost site perturbed.

**Figure S5:**
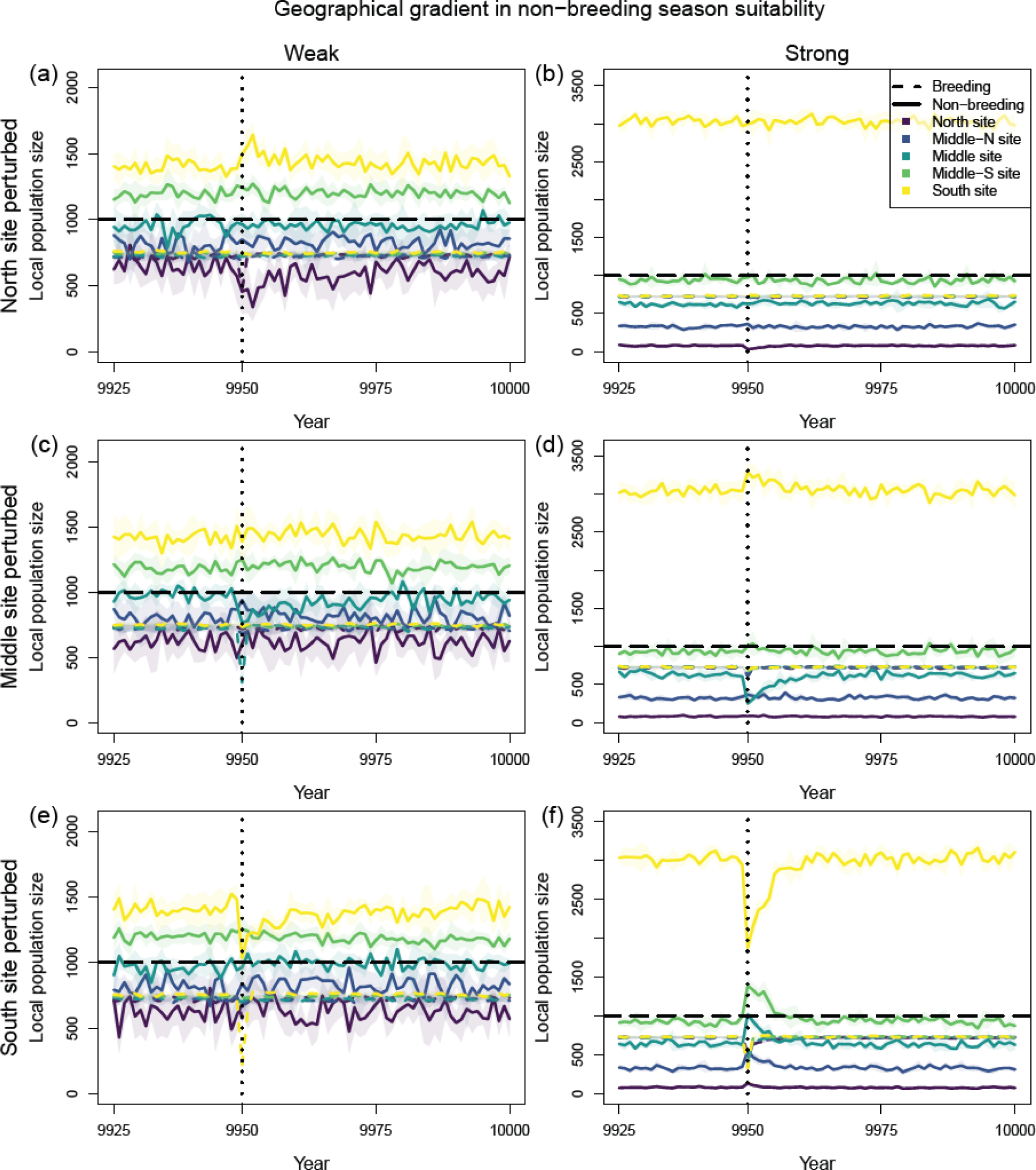
As Fig. S4, except that the metapopulation has evolved when costs of migration depend on the distance between patches travelled.

## Appendix S3: Effects of ECEs when fecundity is lower

In the results presented in the main text, we observed a divergence in time-scales of recovery for population versus evolutionary dynamics following an extreme climatic event (ECE), with breeding population sizes recovering rapidly (2-5 years, see Fig. 5, dashed lines) after a strong local mortality event. Meanwhile, evolutionary responses in migratory frequencies and destinations persisted for multiple generations (10-25 years, see Fig. 5, solid lines, and Fig. 6), causing lasting changes in metapopulation non-breeding season space use. Here we examine whether these contrasting response times remain even if population intrinsic rates of increase are reduced, by changing maximum fecundity *f* to 2 rather than 3 (compare Fig. S6 vs. Fig. 5; Fig. S7 vs. Fig. 6).

Indeed, the same divergence in response time does persist with *f*=2. Breeding population sizes still recover quickly (<5 years, dashed lines in Fig. S6), whereas non-breeding season population distributions are determined by evolutionary changes (effects i-iii described in main text and Appendix S2) and thus recover more slowly (still 10-25 years, solid lines in Fig. S6). The weaker effect of all concurrent evolutionary changes, as well as their slower recovery (due to less overproduction of offspring and a larger component of drift), is evident in all three effects (i-iii) when comparing Fig. S7 and Fig. 6:

i. The decrease in breeding values for migration in the subpopulation breeding at the perturbed site is smaller in Fig. S7 and its recovery is slower.
ii. The increases in breeding values in all other subpopulations are smaller in Fig. S7 and their recoveries slower (up to 30 years).
iii. The shifts in the Destination loci away from the perturbed (South) site are smaller in Fig. S7, and their recoveries slower (up to 30 years).

**Figure S6:**
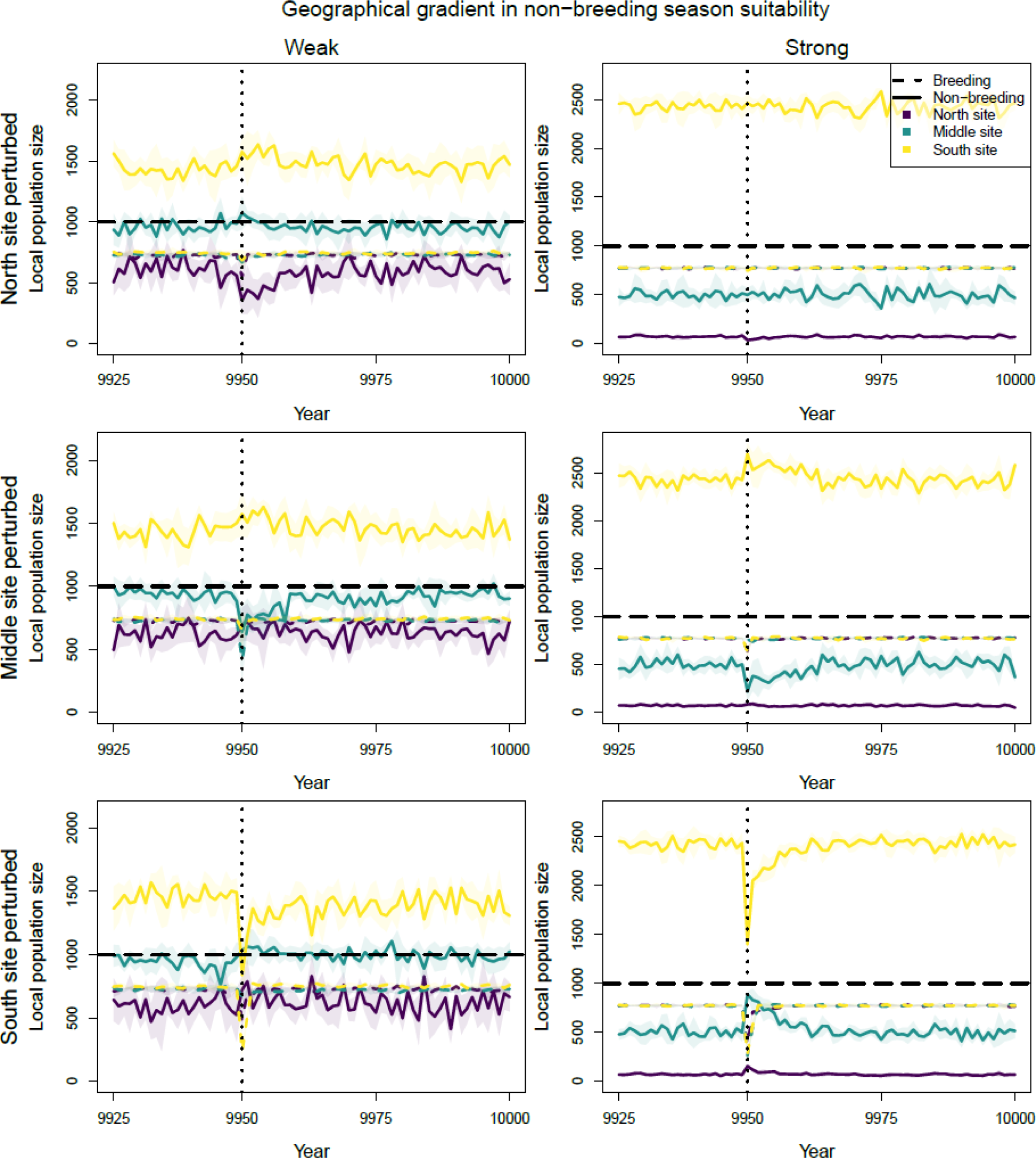
Population dynamics of evolved partially migratory metapopulations before and after an extreme perturbation during the non-breeding season of year 9950 (vertical dotted lines), when maximum fecundity *f*=2 rather than 3. Breeding (dashed colored lines) and non-breeding season (solid colored lines) local population densities are shown for each site. Thick colored lines and opaque bands represent means and 95% CIs across replicate simulations. Dashed black line represents carrying capacity. (a, b): North site perturbed. (c, d): Middle site perturbed. (e, f): South site perturbed. (a, c, e) Weak geographical gradient in non-breeding season suitability (density-dependent mortality parameter ***γ*_n_** = {1/2, 1/3, 1/5}). (b, d, f) Strong geographical gradient in non-breeding season suitability (***γ*_n_** = {5, 1/2, 1/10}). Other parameters set to baseline values (see Appendix S1: Table S1).

**Figure S7:**
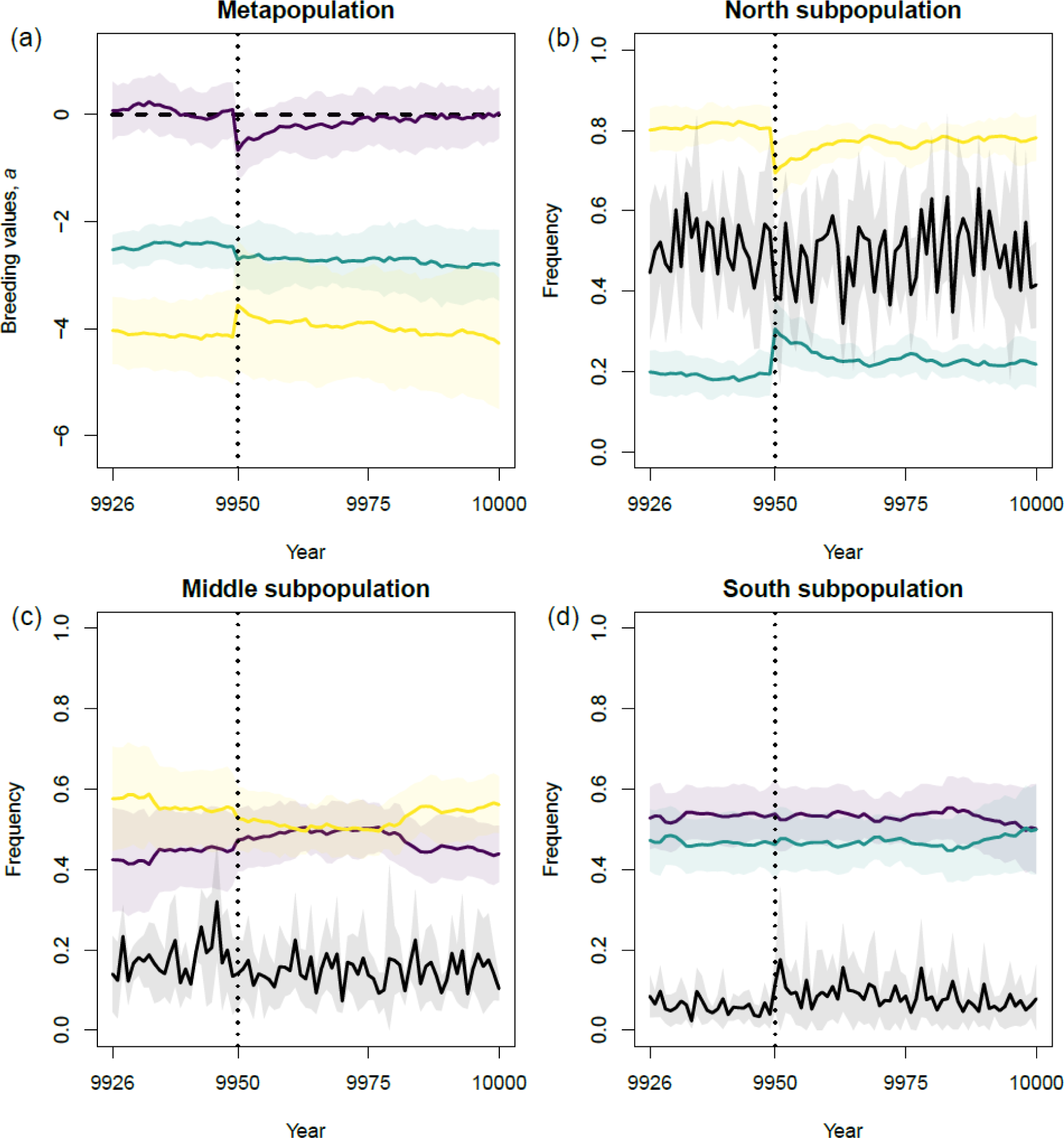
Evolutionary consequences of an extreme climatic event (ECE) striking the south site during the non-breeding season in year 9950 (vertical dotted lines), as in Fig. 5e, Fig. S6e, except maximum fecundity *f*=2 rather than 3. Weak gradient of non-breeding season suitability, ***γ*_n_** = {1/2, 1/3, 1/5}. (a) Subpopulation mean breeding values across replicate simulations (opaque bands show 95% CI) before and after extreme event. Purple, green and yellow show respectively north, middle and south subpopulation. (b)-(d): For each subpopulation (panel header), colored lines show allele frequencies at destination locus (purple, green and yellow show respectively north, middle and south destination allele), and black lines migrant frequencies. Lines show means and opaque bands 95% CI across replicate simulations.

